# Multiscale 3D Genome Reorganization during Skeletal Muscle Stem Cell Lineage Progression and Muscle Aging

**DOI:** 10.1101/2021.12.20.473464

**Authors:** Yu Zhao, Yingzhe Ding, Liangqiang He, Yuying Li, Xiaona Chen, Hao Sun, Huating Wang

## Abstract

3D genome rewiring is known to influence spatiotemporal expression of lineage-specific genes and cell fate transition during stem cell differentiation and aging processes. Yet it is unknown how 3D architecture remodels and orchestrates transcriptional changes during skeletal muscle stem cell (also called satellite cell, SC) activation, proliferation and differentiation course. Here, using *in situ* Hi-C we comprehensively map the 3D genome topology reorganization at multiscale levels during mouse SC lineage progression and integrate with transcriptional and chromatin signatures to elucidate how 3D genome rewiring dictates gene expression program. Specifically, rewiring at compartment level is most pronounced when SC becomes activated. Striking loss in TAD border insulation and chromatin looping also occurs during early activation process. Meanwhile, TADs can also form TAD clusters and super-enhancer containing TAD clusters orchestrate stage-specific gene expression during SC early activation. Furthermore, we elucidate 3D chromatin regulation of key transcription factor, PAX7 and identify cis-regulatory elements that are crucial for local chromatin architecture and *Pax7* expression. Lastly, 3D genome remodeling is profiled in SCs isolated from naturally aging mice, unveiling that geriatric SCs display a prominent gain in long-range contacts and loss of TAD border insulation. Genome compartmentalization and chromatin looping are evidently altered in aged SC while geriatric SC display a more prominent loss in strength of TAD borders. Together, our results implicate 3D chromatin extensively reorganizes at multiple architectural levels and underpin the transcriptome remodeling during SC lineage development and SC aging.

## Introduction

Skeletal muscle tissue has a remarkable regenerative capacity, which largely depends on the activation of resident muscle stem cells, named satellite cells (SCs)^1^. These cells normally reside in a niche beneath the basal lamina of the myofibers in a quiescent state and uniquely marked by the expression of paired box 7 (Pax7) protein. Upon injury, SCs rapidly activate to enter into cell cycle, undergo proliferative expansion, differentiate and eventually fuse to form multinucleated myotube cells; these myotubes further mature into myofibers to repair the damaged muscle. A subset of SCs undergo self-renewal and return to the quiescent state to replenish the adult stem cell pool^2^. Deregulated SC activity is linked to the development of many muscle associated diseases, which necessitates the comprehensive understanding of the mechanisms governing SC fate transition^3–5^. Gene regulation is the key to understand the molecular mechanisms governing SC function. For example, it is well known that *Pax7* is uniquely expressed in SCs and plays multifaceted roles in regulating SC quiescence, proliferation, and self-renewal^6^. PAX7 protein is abundantly enriched in quiescent SCs but gradually diminishes as cells become activated, proliferating and committed to terminal differentiation. However, the regulatory mechanisms underpinning the temporal expression of *Pax7* are largely elusive.

Recent advents in mapping genome-wide 3D chromosomal interactions have led to unprecedented understanding of hierarchical organization of the genome within the 3D nucleus and how gene expression is orchestrated at 3D level^7–11^. At intermediate scales (200 kb to 1 Mb), TADs are identified from chromosome contact maps, which correspond to sequences that interact with each other more frequently within the same TAD rather than with sequences outside the TAD. In mammalians, TAD borders are often demarcated by CTCF and cohesin complex; perturbation of TAD borders is emerging as a driver of many human diseases^12–14^. At a larger scale (multi-Mb), interactions between TADs are thought to form two main types of compartments termed A and B compartments, which are associated with distant chromatin modifications and gene expression. In contrast to TADs which are largely conserved among different cell types and species, compartments can display high variability. Recent studies also reveal that non-contiguous TAD clusters can act in concert to organize genome topology^15–17^. Whereas long-range TAD interactions are enriched in B compartments and correlate with gene repression^15^, super-enhancer containing and highly transcribed TADs can also form clusters that span tens of megabases^16^. When high-resolution Hi-C data is available, chromatin loops can be further identified in Hi-C contact matrix. In general, two types of chromatin loops are revealed: one is constitutive and demarcates TAD borders; the other is more cell-type specific and is often associated with enhancers. Recent findings suggest the 3D genome is commonly reorganized during stem cell lineage progression and differentiation to ensure the spatiotemporal expression of lineage-specific genes^7, 10, 18, 19^. For example, during mouse ESC differentiation into cortical neurons, interactions between B compartments increase with a concomitant decrease of interactions within A compartments; promoter-enhancer contacts are established concomitantly with changes in gene expression^7^. Despite the rapidly gained understanding of 3D genome rewiring during stem cell differentiation, the knowledge gaps remain to be filled in SC. In particular, our knowledge on the molecular events governing early steps of quiescence maintenance and activation of SC is scarce because of the difficulty in preserving SCs in its native niche environment^20, 21^. In fact, the lengthy dissociation steps expose cells to stress and disrupt the physiological niche, which alters SC transcriptome and histone modifications inevitably; fixation prior to isolation has been shown to preserve the quiescent status of SCs^20, 22^, making it possible to investigate whether the 3D genome organization remodels in the early stages of SC activities and whether the 3D rewiring dictates transcriptional changes.

Alterations at 3D levels are associated with aging in many tissues and cells^23–26^. Skeletal muscle suffers from sarcopenia, i.e. functional and mass decline, during human aging; this can partly be attributed to the decrease in the number and function of SCs^1, 4, 5, 27, 28^. Elucidating the molecular causes underlying SC aging thus has become the key to better understand sarcopenia and uncover novel approaches for managing and treating muscle weakening conditions during physiological aging. Research from recent years demonstrates that the SC dysfunction during aging process can be ascribed to changes in transcriptional networks and chromatin state^27, 29–31^. Yet, it is still unknown whether 3D deregulation occurs in aging SC and drives the aging process.

Here, we comprehensively map the 3D genome reorganization during SC lineage progression. We reveal the 3D chromatin extensively reorganizes at all scales including compartment, TAD, TAD cluster and chromatin looping levels. Our results demonstrate dynamic chromatin rewiring at different genomic scales, which orchestrates the transcriptome remodeling and SC activities. Furthermore, in depth analysis at *Pax7* locus uncovers entangling enhancer-promoter connections that establish a unique *Pax7* residing TAD and coordinate its expression dynamics during SC differentiation. Deletion of these regulatory elements disrupts 3D genome organization at *Pax7* locus and attenuates *Pax7* expression. Lastly, we map 3D remolding in SCs isolated from physiologically aging mice and uncover that 3D genome rewiring occurs in aging SCs. Altogether, our findings provide a comprehensive view of 3D genome remodeling during SC lineage progression and SC aging.

## Results

### Global reorganization of genome compartmentalization during SC lineage development

To delineate the remodeling in genome architecture during SC lineage progression, SCs at various stages were prepared (Fig. 1a and Suppl. Fig. 1a). Freshly isolated SCs (FISCs), were collected from Pax7-nGFP mice^32^ by fluorescence-activated cell sorting (FACS) sorting as SCs were exclusively marked by nuclear GFP signal in these mice; based on prior report^20^, these cells were early activating due to the disruption of their niche by the isolation process. To obtain quiescent SCs (QSCs), *in situ* fixation by paraformaldehyde was performed before FACS isolation. Immunofluorescence (IF) staining for PAX7 protein confirmed the purity of the FISCs (91% ± 4.7% positive for PAX7) and QSCs (94% ± 8.7% positive for PAX7) (Suppl. Fig. 1b). FISCs were then cultured *in vitro* for 1, 2 or 3 days to become activated SCs (ASC-D1), proliferating (ASC-D2) or early differentiating (DSC-D3) cells (Fig. 1a). 3D genome organization in the above cells was then probed by *in situ* Hi-C with two or three biological replicates per sample (Fig. 1a and Suppl. Fig. 1c). Following the standard pipeline of data analysis using HiC-Pro^33^, a total of over ∼2 billion uniquely aligned contacts were obtained from all samples (Suppl. Data 1). The first principal component (PC1) values of the biological replicates of each time point were highly correlated (R ≥ 0.95) (Suppl. Fig. 1c); and a gradual decrease in PC1 correlation was observed (Suppl. Fig. 1d) as cells progressed from QSC to DSC-D3, suggesting changes in 3D genome organization were highly reproducible for each time point and cumulative during the SC lineage development. The data from the above biological replicates were thus merged for calculation of *cis*/*trans* interaction ratios; as a result, over 70% of *cis* interactions were observed across all time points (Suppl. Fig. 1e, Suppl. Data 1), reinforcing the high quality of libraries^34^. A resolution of 5 kb for QSC and FISC, and 15 kb for D1, D2 and D3 were obtained after in-depth sequencing, which rendered us to explore the 3D genome architecture at multi-scales, including compartments, TADs and chromatin loops (Fig. 1a).

**Fig. 1.**
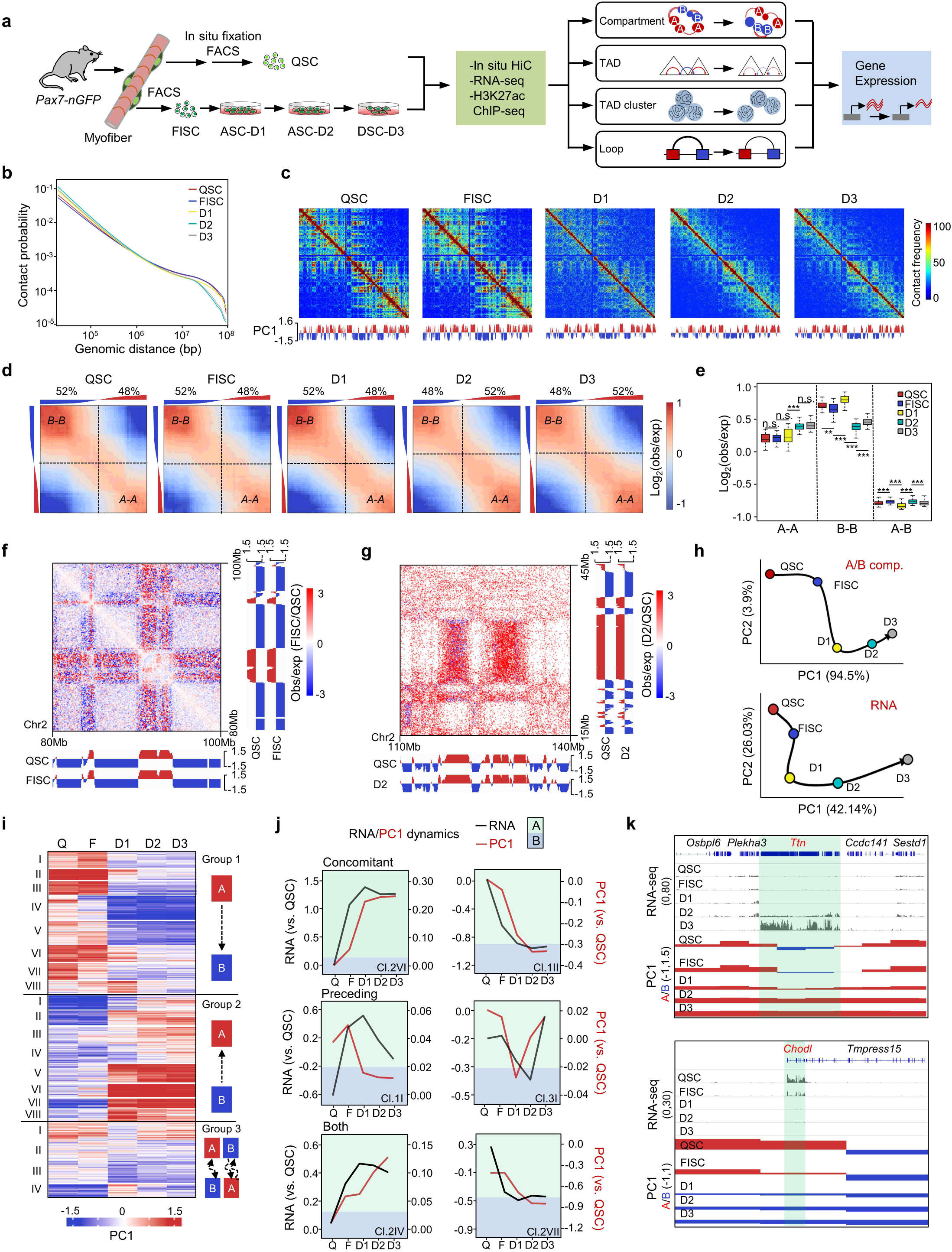
Global reorganization of genome compartmentalization during SC lineage development. **a** Schematic overview of the study design. SCs were isolated from 2 month old Pax7-nGFP mouse by FACS sorting with or without the *in situ* fixation to obtain quiescent (QSCs) or freshly isolated SCs (FISCs) respectively. FISCs were cultured in vitro for various time (D1, D2 and D3) to collect activated (ASC) or differentiating (DSC) SCs. The collected cells were subject to *in situ* Hi-C and RNA-seq. RNA-seq data from QSC and H3K27Ac ChIP-seq data from QSC and FISC were obtained from Léo Machado et. al.^20^ The above datasets were integrated to elucidate 3D genome remodeling at compartment, TAD, TAD cluster, and chromatin loop levels and how 3D rewriting instructs gene expression dynamics during the SC lineage progression. **b** Contact probability was calculated as a function of genomic distance for interactions within individual chromosome across the whole genome at each stage. **c** Observed contact matrices for chr1 at 500-kb resolution and the first eigenvector at 100-kb resolution during the course of SC lineage progression. **d** Saddle plots showing the genome-wide compartmentalization over time. Percentage of compartment A/B and PC1 value of each 100-kb loci arranged by eigenvector are shown on top. **e** Contact enrichment of 100-kb loci between compartments of the same (A vs A or B vs B) and different (A vs B) types. For all box plots, center lines denote medians; box limits denote upper and lower quartiles; whiskers denote 1.5× interquartile range. P-values were calculated by a Wilcoxon signed-rank test. **f** Differential heatmap for 20Mb regions on chromosome 2 showing increased A-B interactions in FISC vs QSC. PC1 signal at each stage is displayed on the bottom of the heatmap. **g** Differential heatmap for a 30 Mb region on chromosome 2 showing an increased A-A interaction in FISC vs. QSC. **h** PCA of PC1 values and gene expression dynamics during SC lineage progression. Black arrows denote the hypothetical trajectory. **i** *k*-means clustering (*k* = 20) of PC1 values for 100-kb genomic bins that switch compartment at any time point during SC lineage progression. **j** Examples of individual switching clusters with concomitant PC1 and gene expression changes (VI/group2), and clusters with PC1 changes preceding expression changes (II/group1), clusters with expression changes preceding PC1 changes (I/group, I/group3) or clusters with both IV/group2 and VII/group2 occurring. **k** Genome-browser view of *Osbpl6-Ttn-Sestd1* (top) and *Chodl-Tmpress15* (bottom) loci. PC1 and RNA-seq values at each stage are shown. Green shadings denote B-A-switch (top) and A-B-switch (bottom) regions.

Overall, comparing the later (D2 and D3) with the early (QSC, FISC and D1) stages we observed a dramatic decrease in the frequency of long-range (> 2 Mb) but an increase in the short-range (< 2 Mb) interactions on all chromosomes (Fig. 1b, c and Suppl. Fig. 1f), implying that from a genome-wide perspective, the 3D genome organization underwent distinct remodeling in the later stages when cells started to proliferate. Genome segregation into A and B compartments was then determined (Suppl. Data 2). As expected, A-type compartments were associated with active histone marks, H3K4me3 and H3K27ac (ChIP-seq data were obtained from Léo Machado *et. al.*^20^), which were not present in B-type compartments (Suppl. Fig. 1g). Although the overall proportions assigned to A (47%-50%) and B (50%-53%) compartments remained stable throughout the entire lineage progression (Suppl. Fig. 1h), compartmentalization strength (as measured by the average contact enrichment within (A-A, B-B) and between (A-B) compartments) was dynamically altered. A more condensed chromatin state of compartment B was observed at earlier stages (QSC, FISC and D1) with a larger proportion of B-B interactions compared to later stages (D2 and D3) (52% vs 48%) (Fig. 1d) and a significantly higher interaction strength (Fig. 1e); the proportion of A-A interactions and interaction strength, on the contrary, markedly increased at later stages (Fig. 1d-e). For instance, a loss of compartmentalization was observed in FISC vs. QSC as revealed by reduced B-B contact frequencies and increased inter-compartment A-B contacts (Fig. 1f), but recovered A-B compartment segregation was observed at D2 vs. QSC (Fig. 1g). Switching of loci between A and B compartments was examined and only 12% of the genome switched compartment at any two time points with B to A and A to B each accounting for 6% (Suppl. Fig. 1i); interestingly, 32% of these regions were involved in multiple switching events (Suppl. Fig. 1i) and the switched genomic regions exhibited the most substantial PC1 changes (Suppl. Fig. 1j), which is consistent with previous findings^9^ .

Next, harnessing the existing transcriptomic profiles from QSC^20^ and the in house generated RNA-seq from FISC, D1, D2 and D3 cells (Suppl. Data 3), the interplay between compartmentalization and gene expression changes was examined (Suppl. Data 3). Consistent with the lineage progression timing, we observed high expression of *Pax7* in QSCs but its level sharply declined in early activating FISCs; *MyoD* was on the other hand absent in QSCs but highly induced in FISCs and maintained the expression at later stages; *MyoG* was only induced when SCs initiated differentiation at D3. *HeyL* was decreased while *Egr1* was induced in FISC vs QSCs (Suppl. Fig. 2a). Overall, the trajectory of genome compartmentalization demonstrated a drastic change occurred at the transition of QSC to FISC, which was not observed on the transcriptomic trajectory (Fig. 1h). *k*-means clustering was then performed based on the PC1 values of the 20% of the genome (n = 2,605 genes) with compartment switching (Suppl. Data 2). A total of 20 clusters with a broad range of switching dynamics were identified (Fig. 1i, Suppl. Fig. 2b); 6 of them (1II, 1III, 1VII, 2VI, 2VIII, and 3IV) demonstrated concomitant changes in compartmentalization and gene expression (R > 0.8; Fig. 1j), but this correlation was lost at least at one time point in other clusters (Supp. Fig. 2b).

Specifically, 7 out of the 20 clusters displayed compartment changes preceding transcriptional changes, and these clusters involved half of the genes with switched compartment (Suppl. Fig. 2b, cluster 1I); only two clusters displayed compartment changes lagging behind gene expression changes whereas 5 clusters displayed both preceding and lagging relationships. For example, as shown in Fig. 1k (top), *Ttn* locus which encodes a contractile and sarcomere structural protein ^35, 36^, was embedded in B compartment in QSC and switched from B to A at D1 stage which was further strengthened at D3; transcription activation of the *Ttn*, on the other hand, was initiated at D2 and strongly enhanced at D3. In another example, *Chodl* (Fig. 1k, bottom), recently identified as a QSC-signature gene^20^, showed a sharp decrease of its expression in FISC vs. QSC and completely undetectable at later stages (D1-D3). Accordingly, we noticed a gradual transition from A to B compartment at *Chodl* locus from QSC to FISC; and B-compartment association was enhanced at D1-D3 stages. Furthermore, Gene ontology (GO) analyses demonstrated that genes in all the above 20 clusters undergoing compartment switch were associated with metabolic (negative regulation of DNA demethylation, regulation of the apoptotic process, protein auto phosphorylation etc.), secretory functions (extracellular exosome etc.) and developmental processes (cell division& cell cycle, positive regulation of fibroblast proliferation, cell development, cell differentiation etc.) (Suppl. Fig. 2c and Suppl. Data 3). Altogether, the above findings demonstrate that genome compartmentalization is dynamically rewired during SC lineage progression and most pronounced when cells become activated; and compartmentation remodeling is tightly coupled with the transcriptomic reprogramming with compartmentalization changes preceding expression changes in a substantial number of genes.

### Dynamic TAD organization during SC lineage development

Next, we examined TAD organization during SC lineage development. First, an average of ∼3,000–3,400 borders per time point was identified by using Topdom^37^ (Fig. 2a and Suppl. Data 4). Partitioning of the genome into TADs was largely stable during the lineage progression as >60% TAD borders (n = 2,155) were detected at all stages (Fig. 2a, invariant borders); a comparable number of TAD borders were stably acquired (704/4,892, 14.4%) or lost (779/4,892, 15.9%), thus resulting in a steady number of borders and conserved average TAD size between 720 to 760 kb (Suppl. Fig. 3a). Nevertheless, border reorganization was observed at the transition of each two stages. For example, 535, 95, 116, 66 and 179 TADs rearranged, merged, split, built, or disappeared at the transition from FISC to D1 (Suppl. Fig. 3b). An insulation score (IS) was then calculated to quantitatively measure the local chromatin insulation strength by TAD borders (Suppl. Data 4). In line with previous reports^9^, the number of CTCF binding motif was negatively correlated with the IS for all borders (Suppl. Fig. 3c), indicating CTCF occupancy increased the insulation strength of TAD borders. Quantitative measurement of the borders that were gained (Suppl. Fig. 3d) or lost (Suppl. Fig. 3e) in each stage relative to QSC revealed that borders gained in later stages (D1-D3) displayed an IS decrease in the early stages (FISC and D1) (Suppl. Fig. 3d). Similarly, borders lost in the later stages already exhibited considerable increase of IS in the beginning stages (Suppl. Fig. 3e). Aggregate interaction matrix centered on borders of each stage revealed a genuine loss of insulation across the TAD borders early in FISC vs. QSC (Fig. 2b). IS also increased drastically in FISC vs. QSC but decreased when SCs activated, proliferated and differentiated (D1, D2, D3 vs. FISC) (Fig. 2c). Furthermore, compared with qualitative border changes, quantitative changes in IS of the invariant TADs were more evident: hierarchical clustering identified that 52.8% (1,133/2,155) of all invariant borders showed > 50% difference in between each two time points (Fig. 2d). Altogether, the above findings demonstrate that border re-organization gradually occurs during the entire lineage development. Although distinct TAD re-organization is observed in late vs. early activation stages (D1 vs FISC), the most substantial changes in TAD border strength occur in early activation stages (FISC).

**Fig. 2.**
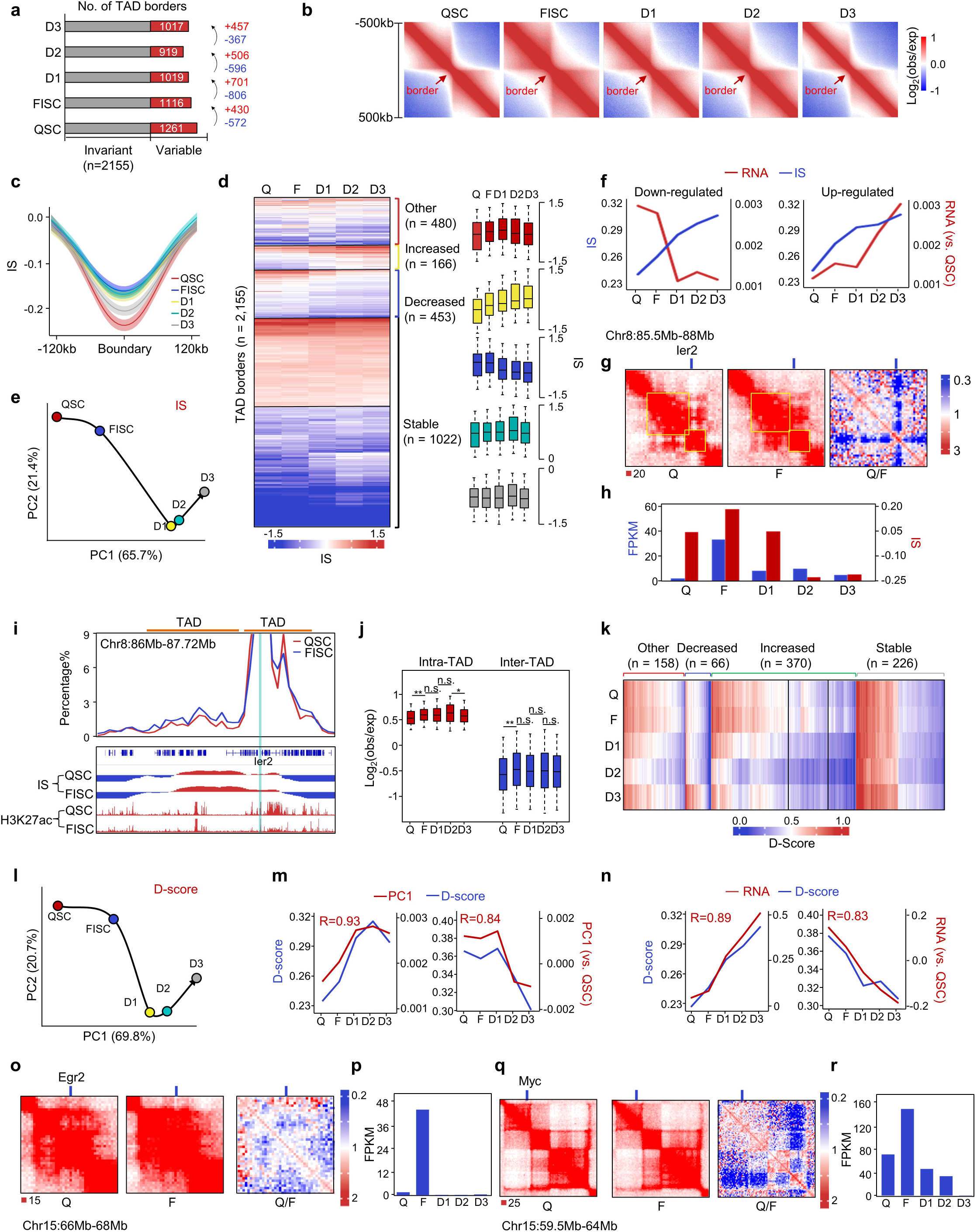
Dynamic TAD organization during SC lineage development. **a** The number of TAD borders identified per time point. Invariant borders are present at all stages while variable borders are lost or acquired in between two adjacent stages. **b** Aggregate Hi-C maps in a 1Mb region centered on the invariant TADs boundaries. Data are presented as the log ratio of observed and expected contacts in 40kb bins. **c** Average insulation score (IS) in a 240 kb region centered on invariant TADs boundaries. Lines denote the mean values, while the shaded ribbons represent the 95% confidence interval (CI). **d** *k*-means clustering (*k* = 20) of IS. Bar graphs show kinetics for groups that increased (n = 1,258), decreased (n = 527), transiently increased (n = 254) or did not change (n = 2,632) IS during the lineage progression. **e** PCA of IS during SC lineage progression. Hypothetical trajectory is shown as black arrow. **f** Kinetics of mean expression level (FPKM) changes at dynamic borders harboring up- or down-regulated genes with IS increased. **g** Illustration of *in situ* Hi-C contact maps (40-kb resolution) on the *Ier2* locus. **h** IS kinetics of Ier2 residing TAD border and its expression dynamics. **i** (top) Representative virtual 4C analysis on *Ier2* locus in QSC and FISC. Ier2 residing and upstream TADs are indicated. (bottom) Genome-browser view of IS and H3K27ac ChIP-seq are shown. **j** Contact enrichment of intra- and inter-TAD during SC lineage progression. Data is represented as a boxplot based on the intra- and inter-TAD values per TAD. For all box plots, center lines denote medians; box limits denote upper and lower quartiles; whiskers denote 1.5× interquartile range. P-values were calculated by a Wilcoxon signed-rank test. **k** *k*-means clustering (*k* = 20) of D scores. Bar graphs show D-score kinetics for groups that increased (n = 2,172), decreased (n = 467), or stable (n = 1,456). **l** PCA of D-score during SC lineage progression. Hypothetical trajectory is shown as black arrow. **m, n** Average D-score and PC1 kinetics (**m**) or gene expression (**n**) analysis for clusters of TADs that were gained (n = 2,172, left) or lost (n = 467, right) in D3 vs QSC. Pearson correlation coefficients (R) are indicated. **o** Illustration of in situ Hi-C contact maps (40-kb resolution) on the Egr2 locus on Chr15. **p** The bar graphs show the FPKM kinetics of Egr2 expression. **q** Illustration of in situ Hi-C contact maps (40-kb resolution) on the Myc locus on Chr15. **r** The bar graphs show the FPKM kinetics of Myc expression.

To further investigate if TAD reprogramming is correlated with the local gene expression, we found that PCA of IS kinetics (Fig. 2e) showed a progressing trajectory grossly resembling the transcriptome and PC1 trajectories determined earlier (Fig. 1h) despite the qualitative gain or loss of TAD borders did not correlate with overall increased (Suppl. Fig. 3f) or decreased (Suppl. Fig. 3g) changes of local gene expression. When examining transcriptional changes at the most dynamic 545 borders that increased (Fig. 2f) or decreased (Suppl. Fig. 3h) in quantitative IS measurement (|(*IS_D_*_3_ − *IS_QSC_*)/ *IS_QSC_*| ≥ 2), the majority (81.8%, 446/545) were associated with local gene down-regulation (n = 548) or up-regulation (n = 41) (log_2_FC> 1.5 between endpoints) (Fig. 2f, Suppl. Fig. 3h). Unexpectedly, IS increase did not appear to proceed the transcriptional changes, instead concomitant changes were observed on many loci. For example, in QSC, the TAD surrounding *Ier2* locus was insulated by a strong border with a low IS; the IS considerably increased in FISC resulting in a merge with an upstream TAD (Fig. 2g) and the transcriptional level increased concomitantly (Fig. 2h). Consistently, Hi-C-derived virtual 4C also revealed the increased interaction between the TAD of *Ier2* with the upstream one in FISC vs. QSC (23.4% vs. 16.3%) (Fig. 2i). As further illustrated on *HeyL* locus (Suppl. Fig. 3i), the associated TAD border weakened with increased IS (Suppl. Fig. 3i), which might result in the concomitant decrease of transcriptional level from QSC to FISC (Suppl. Fig 3j).

Next, to determine the self-contacting propensity within a given TAD, we calculated a domain score (D-score) to measure the portion of intra-TAD interactions in all *cis* interactions (Suppl. Data 4). As previously noted^9^ D-scores were positively associated with gene expression (Suppl. Fig. 3k). Hierarchical clustering revealed that 72% (n = 594) of the invariant TADs displayed D-score change (> 20% difference between each two time points) (Fig. 2k). Additionally, PCA of D-score kinetics revealed a trajectory (Fig. 2l) resembling those of compartmentalization, transcription, and IS. Overall, The D-score kinetics correlated closely with compartmentalization (PC1) changes (R > 0.84; Fig. 2m). Similar to IS, it was also highly correlated with changes in gene expression (R >0.83, Fig. 2n). For example, the D score of the TAD associated with *Egr2* locus increased in FISC vs. QSC (Fig. 2o), which led to concomitant transcriptional activation of *Egr2* (Fig. 2p). On *Myc* locus, on the other hand, the D score decrease was accompanied by a transcriptional repression in FISC vs. QSC (Fig. 2q-r). Altogether the above results suggest that the 3D genome is dynamically rewired at TAD level and impinges on gene expression during SC lineage development.

### TADs clusters are dynamically reorganized in FISC vs. QSC

Increasing evidence underscores the importance of large-scale inter-TAD associations that position chromatin domains in the nucleus^19, 38–40^. For example, Paulsen, J. *et. al*^15^ reported a level of developmental genome organization that involves long-range TAD-TAD interactions into assemblies of linearly non-contiguous TADs defined as TAD cliques. Therefore, to further explore the possible organization of inter-TAD interactions during SC lineage progress, we examined our Hi-C contact matrices and indeed uncovered that TADs engaged in non-contiguous contacts (Fig. 3a, exemplified on chromosome 9 at D1 stage). Utilizing non-central hypergeometric model (NCHG), 3639, 3667, 2927, 1843 and 2176 significant intra-chromosomal TAD-TAD interactions were identified from QSC to D3 (Suppl. Data 5), which is exemplified on chromosome 9 at D1 stage (Fig. 3b); the interaction distance between TADs remained constant from QSC to D1 (QSC: 28.6 Mb; FISC: 28.8 Mb; D1: 29.2 Mb) but a decrease was observed starting D2 (19.8 Mb) (Fig. 3c). Next, we defined TAD clusters, which encompassed at least 3 TADs that were fully connected pairwise in Hi-C data regardless of interaction distance (Fig. 3d). As a result, a large number of TADs (QSC to D3: 719, 563, 386, 260, and 284) were engaged in clusters containing 3 to 11 TADs, representing 31-87% of all TADs (Fig. 3e). A reduced number of TADs in the clusters was observed resulting in smaller clusters as cells progressed from QSC to D3 (Fig. 3e). An alluvial diagram showed that whereas 65% of TAD clusters were maintained during the lineage progression, hundreds of TADs assembled into, or disassembled from clusters; waves of TAD cluster formation and dissociation were observed in FISC, D2 and D3, which resulted in variation of cluster size (Suppl. Fig. 4a). Previously, Paulsen, J. et.al.^15^ defined TAD cliques as functionally important TAD assemblies involving long-range TAD-TAD interactions, which are associated with repressed chromatin domain that shapes the genome during cell differentiation and also demonstrated these TAD cliques are enriched in B compartments. Paulsen, J. et. al. ^15^, on the other hand, showed that about 30% of the identified TAD cliques in the adipogenesis residue in compartments A. Interestingly, we found the TAD clusters defined in our study were present in both compartments A and B. For example, a TAD cluster (C1) residing in compartment B in QSC and FISC displayed an increase of cluster size in D1 (4 to 6); a second cluster, C3 assembled at D1 in compartment B, while C2 cluster residing in compartment A underwent dynamic assembly/disassembly during the entire lineage progression (Suppl. Fig. 4b). When closely examining frequency of TAD cluster formation, loss, expansion and reduction at each stage compared to QSC, we found the changes occurred in compartments A and B at about the same frequency in FISC vs. QSC; and similar observation was made in D1 vs. QSC. Nevertheless, changes in D2 or D3 vs. QSC occurred more frequently in compartment B vs. A (Suppl. Fig. 4c). Furthermore, we found that interaction distance between the pairwise TAD-TAD interactions significantly decreased in D1 vs FISC and D2 vs D1 (Suppl. Fig. 4d). Interestingly, TADs in short-range interactions (Q1 and Q2, QSC: <18.26Mb; FISC: <17.98Mb; D1: <18.02Mb; D2: <5.1Mb; D3: <10.24Mb) were nearly equally associated with A and B compartments, while those in long-range interactions (Q3 and Q4, QSC: >18.26Mb; FISC: >17.98Mb; D1: >18.02Mb; D2: >5.1Mb; D3: >10.24Mb) displayed a prominent enrichment towards B compartments (Fig. 3f), suggesting these long-range interacting TADs in Q3 and Q4 were reminiscent of the TAD cliques defined in the previous study^15^. Next, we examined the gene expression in TAD clusters and found genes in Q1 or Q2 were expressed at significantly higher levels than those in Q3 or Q4 or not in any clusters (Fig. 3g), which is consistent with prior findings that TAD clusters involving long-range TAD-TAD interactions are associated with chromatin compaction and gene repression.

**Fig. 3.**
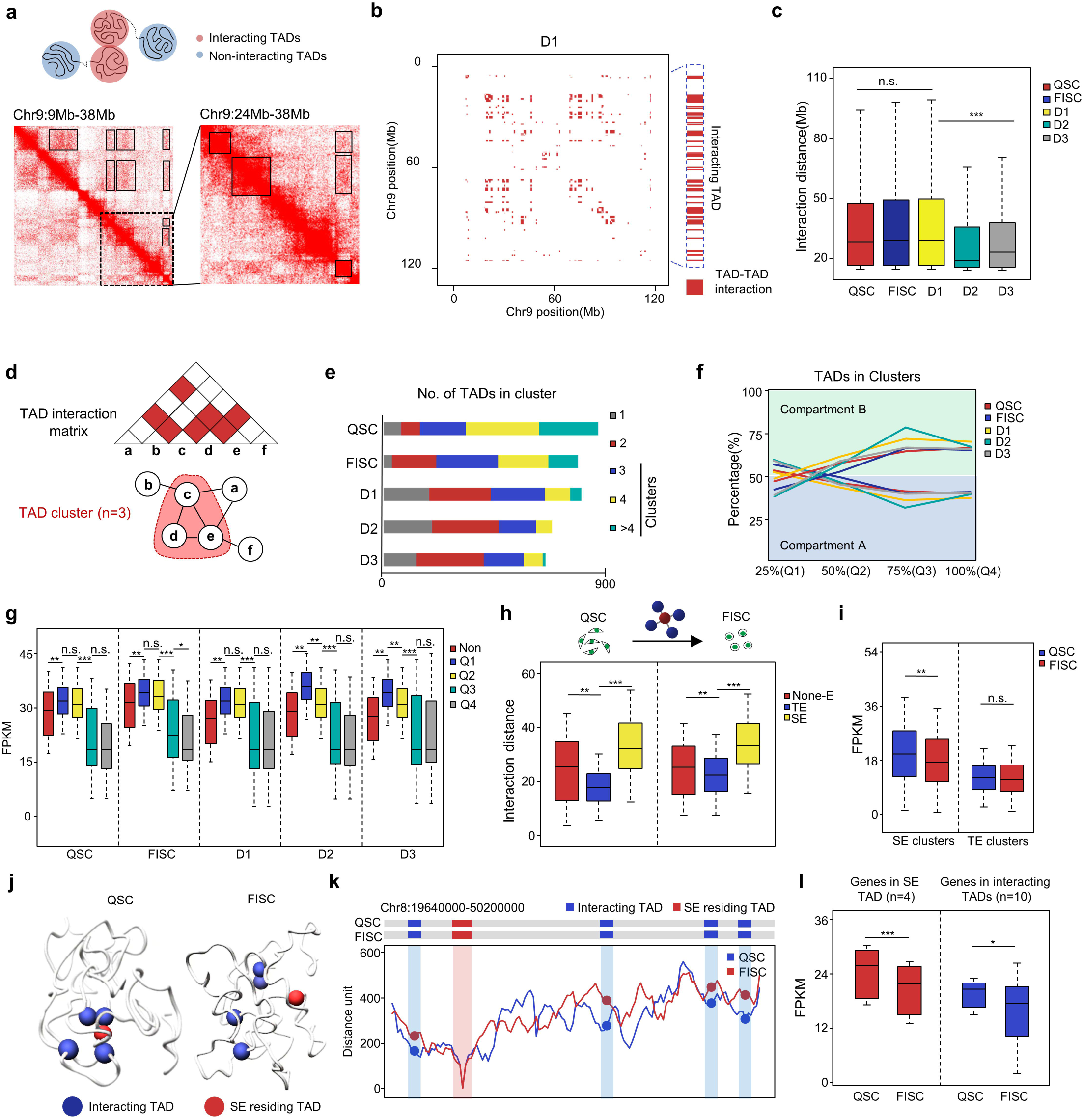
TADs clusters are dynamically reorganized in FISC vs. QSC. **a** (top) Illustration of TAD-TAD interaction (top). (bottom left) a region of TAD-TAD interactions on chromosome 9 (D1) are visualized in the Hi-C data. Boxes, TAD–TAD interactions. the dashed region is enlarged on the right. **b** Matrix of significant TAD-TAD interactions (red pixels) in chromosome 9 (D1); an example of interacting TADs is highlighted (red pixels on the right side); c Boxplots showing the average distance between pair-wise TAD interactions at each time point. Center lines denote medians; box limits denote upper and lower quartiles; whiskers denote 1.5× interquartile range. P-values were calculated by a two-sided Wilcoxon test. **d** (top) illustration of significant interactions between TADs in a matrix (red pixels). (bottom) A TAD cluster (pink) is identified when >=3 TADs are fully connected with each other. Edges indicating interaction between two TADs while nodes indicating TADs. **e** Bar charts depicting the number of TADs in clusters as a function of cluster size at each time point. **f** TADs in clusters were stratified by quartile of distances between pair-wise TAD interactions. Line charts depicting the fraction of TADs in clusters of each quartile in A and B compartments over time of SC development. **g** Gene expression levels outside clusters (Non) and in clusters of each quartile (Q1, Q2, Q3 or Q4) over time of SC development. Box-and-whisker plots show the median FPKM; numbers refer to the number of outliers, including non-protein-coding transcripts. **h** Boxplots showing the average distance between pair-wise TAD interactions that TE or SE engaged in at QSC or FISC stage. **i** Transcription levels in the above SE- or TE-engaged TAD clusters. **j** Top, 3D chromatin conformation model of a SE-engaged TAD cluster identified in QSC and its reorganization in FISC. The SE containing core TAD is marked in red and other interacting TADs are in blue. k line plot at 5 kb resolution displaying the median distance distribution between SE-containing TAD and other interacting TADs in the above cluster at QSC and FISC stages. **l** Expression levels of genes (n=5) in the above SE-containing TAD or in interacting TADs (n = 28). Data in g, h, i and l are presented in boxplots. Center line, median; box limits, upper and lower quartiles; whiskers, 1.5× interquartile range. Statistical analyses were done by Wilcoxon signed-rank test; *P < 0.01, **P < 0.001, ***P < 0.0001.

Lastly, since previous studies revealed that super enhancers (SEs) are enriched in the highly interacting TAD triplets^16, 17, 41^, we wondered whether SEs-containing TAD clusters are more active in long-range TAD-TAD interactions. Leveraging the H3K27ac ChIP-seq profiles in QSC and FISC^20^, we defined SEs and typical enhancers (TEs). Indeed, we found 10%-12% TADs in Q3 and Q4 contained SEs and residing in compartment A (Suppl. Fig. 4e), and those SE containing TAD clusters were associated with higher gene expression in both QSC and FISC compared with non-SE clusters (Suppl. Fig. 4f). Overall, in both QSC and FISC, we found SE-containing TAD clusters were engaged in longer interaction distance (35 kb vs 18 kb) (Fig. 3h), a higher number of TAD clusters (Suppl. Fig. 4g) and a larger cluster size (Suppl. Fig. 4h) compared to those containing TEs or no enhancers. In addition, the presence of SE clusters was associated with more evident expression changes, as transcription level in SE not TE containing TAD clusters in both QSC (Fig. 3i) and FISC (Suppl. Fig. 4i) displayed distinct stage-specific expression pattern. As an example, a 3D model of one TAD cluster was developed (Fig. 3j), which contained one of active SEs in QSC. This SE-containing TAD clustered with four other TADs, to form a TAD cluster in QSC; it decommissioned in FISC due to the increased interacting distance between these TADs (Fig. 3k). Consequently, the genes in this TAD cluster (both the SE residing and four interacting TADs) expressed at significantly lower levels in FISC vs. QSC (Fig. 3l). Taken together, our results demonstrate that TAD clusters are present and dynamically reorganized when SC progresses from QSC to FISC.

### Chromatin loops are dynamically changed during SC early activation

Chromatin loops manifest as foci in high-resolution Hi-C contact maps. Considering the most striking changes observed above at compartment and TAD levels occurred in the early stages (QSC to FISC), we increased the sequence depth for the Hi-C libraries from these two time points to obtain > 500 million uniquely mapped contacts, which enabled us to visualize chromatin looping events at a 5 kb resolution. Using FitHiC2^42^ algorithm, a total of 11,337 and 2412 chromatin loops were identified in QSC and FISC, respectively (Fig. 4a, Suppl. Data 6), indicating a drastic decrease in the number of loops during SC early activation. In addition, a reduction of interaction strength was also observed by aggregated peak analysis (APA) (4.16 vs 3.93) (Fig. 4a). Furthermore, 2,790 and 504 loops were further defined as QSC or FISC specific loops, which accounted for a minor proportion of all loops (24.6% for QSC and 20.1% for FISC) (Suppl. Fig. 5a and Suppl. Data 6). Expectedly, QSC-specific loops showed a decreased APA score in FISC (1.23) vs. QSC (4.12), while FISC-specific loops displayed a much higher APA score in FISC (3.57) vs. QSC (1.91) (Suppl. Fig. 5b). Interestingly, FISC-specific loops were larger than QSC-specific loops and contained fewer number of genes (Suppl. Fig. 5c). However, in both time points the stage-specific loops localized mainly to compartment A (Suppl. Fig. 5d) and positively correlated with gene expression changes (Suppl. Fig. 5e). Moreover, GO analysis revealed genes involved in the stage-specific loops were enriched for distinct terms (Suppl. Fig. 5f and Suppl. Data 6).

**Fig. 4.**
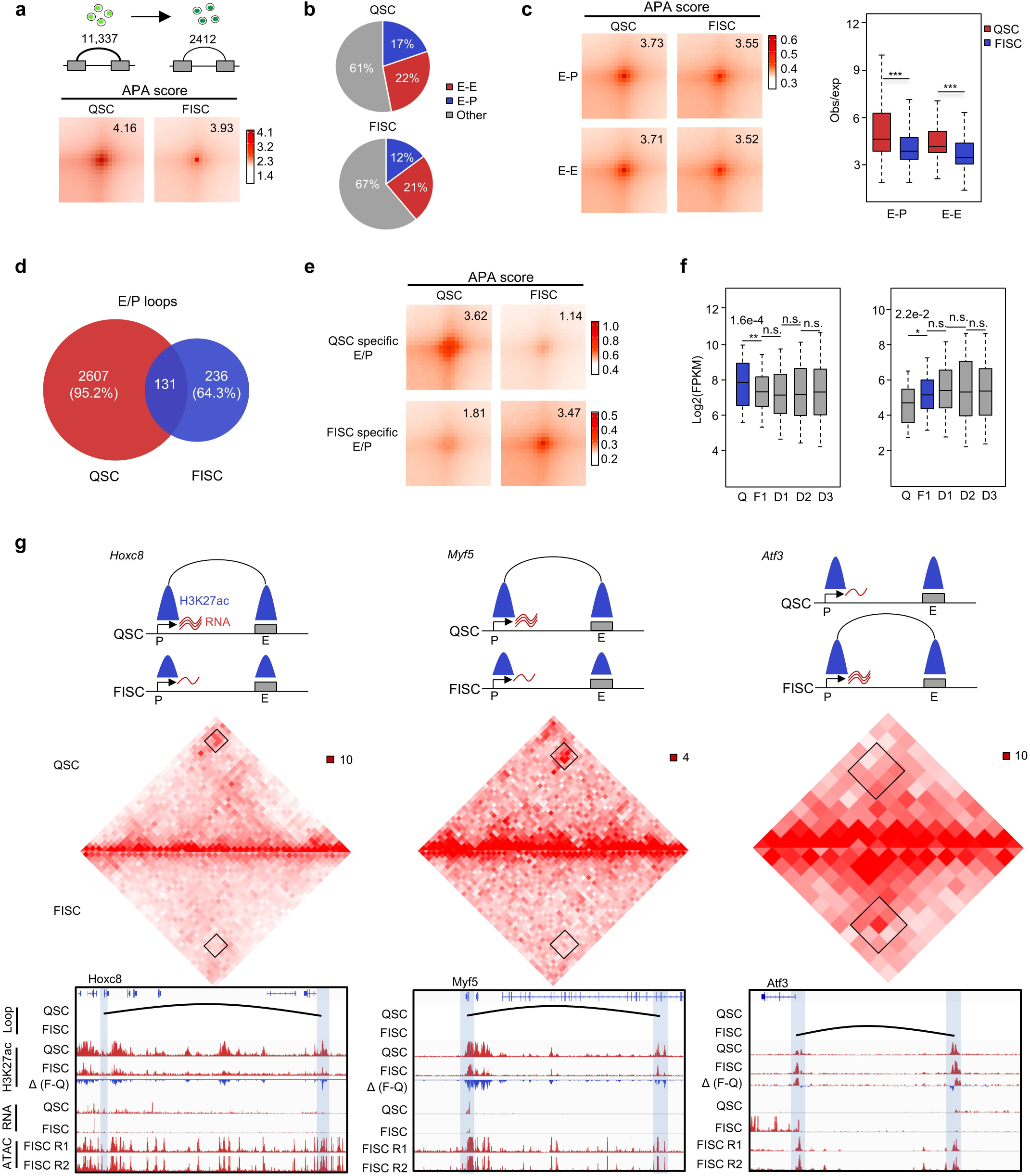
Chromatin loops are dynamically changed during SC early activation. **a** (top) Illustration of identified chromatin loops in QSC and FISC. (Bottom) Aggregated peak analysis (APA) analysis of chromatin loops detected at QSC (n = 2,790) or FISC (n = 504). Bin size, 5 kb. **b** Distribution of enhancer-enhancer (E-E), Enhancer-Promoter(E-P) and other types of loops in QSC and FISC. **c** (left) APA analysis of E-E and E-P chromatin loops detected in QSC and FISC. (right) Box plot depicting interaction intensity (Obs/exp) of the above E-E and E-P loops. **d** Overlapping of E/P loops between QSC and FISC. The number and relative percentage of overlapped and specific E/P loops are shown. **e** APA analysis of interaction intensity of the above QSC or FISC specific E/P loops. **f** Box plot depicting expression dynamics of genes within QSC–specific (left, n = 1,874 genes) or FISC-specific (right, n = 469 genes) loops. **g** (top) Illustration of the dynamics of formation of E-P loop, H3K27ac signal and gene expression on the *Hoxc8* (left), *Myf5* (center) and *Atf3* (right) loci in QSC and FISC. (middle) Hi-C contact maps (5-kb resolution) on the *Hoxc8*, *Myf5*, and *Atf3* loci. (bottom) E-P loops are marked in black curve. Gene promoters and interacting regions are highlighted in blue boxes. Genome browser tracks showing H3K27ac CHIP-seq and RNA-seq profiles in QSC and FISC, ATAC-seq signals of two replicates in FISC are also shown. Data in c and f are presented in boxplots. Center line, median; box limits, upper and lower quartiles; whiskers, 1.5× interquartile range. Statistical analyses were done by Wilcoxon signed-rank test; *P < 0.01, **P < 0.001, ***P < 0.0001.

Next we investigated subtypes of loops formed between enhancers and promoters (E-P) or two enhancers (E-E). Of all E/P related loops in QSC, 17% were found to be E-P while 22% were E-E interactions; in FISC, 12% were E-P while 21% were E-E interactions (Fig. 4b). Interestingly, enhancers in E-E interactions displayed lower levels of H3K27ac signals in FISC vs QSC, while promoters involved in E-P interactions showed higher levels of H3K4me3 signals, indicating that E-E or E-P loops were epigenetically distinct in QSC and FISC (Suppl. Fig. 5g). The interaction strength of both E-P (APA: 3.73 to 3.55) and E-E loops (APA: 3.71 to 3.52) decreased in FISC vs. QSC (Fig. 4c). Furthermore, we found although enhancer/promoter landscapes were largely unaltered in FISC vs QSC (10,113 shared enhancers/promoters) (Suppl. Fig. 5h), the loop formation was highly dynamic with 95.2% and 64.3% specifically identified in QSC and FISC respectively (Fig. 4d). In addition, stage-specific E-P and E-E loops also displayed more prominent difference in APA score (1.14 vs. 3.62) (Fig. 4e) and correlation with gene expression (Fig. 4f), when compared to all loops (Suppl. Fig. 5e). This phenomenon is illustrated on representative *Hoxc8*, *Myf5* and *Atf3* genes (Fig. 4g). For both *Hoxc8* and *Myf5,* a disappeared E-P loop in FISC vs. QSC was observed between the gene promoter and their distal enhancers which displayed the openness at both stages. As a result, the gene expression was decreased in FISC. On the contrary, on *Atf3* locus, despite the openness of the distal enhancer in QSC, the gene expression was not observed until the E-P loop became established in FISC. Taken together, these results reveal dynamic changes of looping events during early activation of SCs, which may be tightly related to the transcriptional change of pivotal genes associated with this progression.

### 3D regulatory interactions orchestrate Pax7 expression dynamics in SC

PAX7 is the key TF known to regulate the myogenic potential and function of SCs^43^, the mechanism dictating its high expression in QSC and attenuation after SC becomes activated remains largely unknown. We thus took a close examination at the 3D organization associated with *Pax7* locus. A prominent TAD was identified in QSC encompassing *Pax7* locus only and evident presence of four sub-TADs was also observed (Fig. 5a). Whereas the TAD maintained constant, the four sub-TADs disappeared in FISC due to a loss of interaction strength within the sub-TADs and a gain of interaction strength among them (Fig. 5a-b). In addition, a reduced number of chromatin loops were observed in FISC (3) vs. QSC (9) (Fig. 5b). The above data suggest that *Pax7* locus undergoes drastic reorganization at 3D level when SCs become activate which may account for its expression attenuation.

**Fig. 5.**
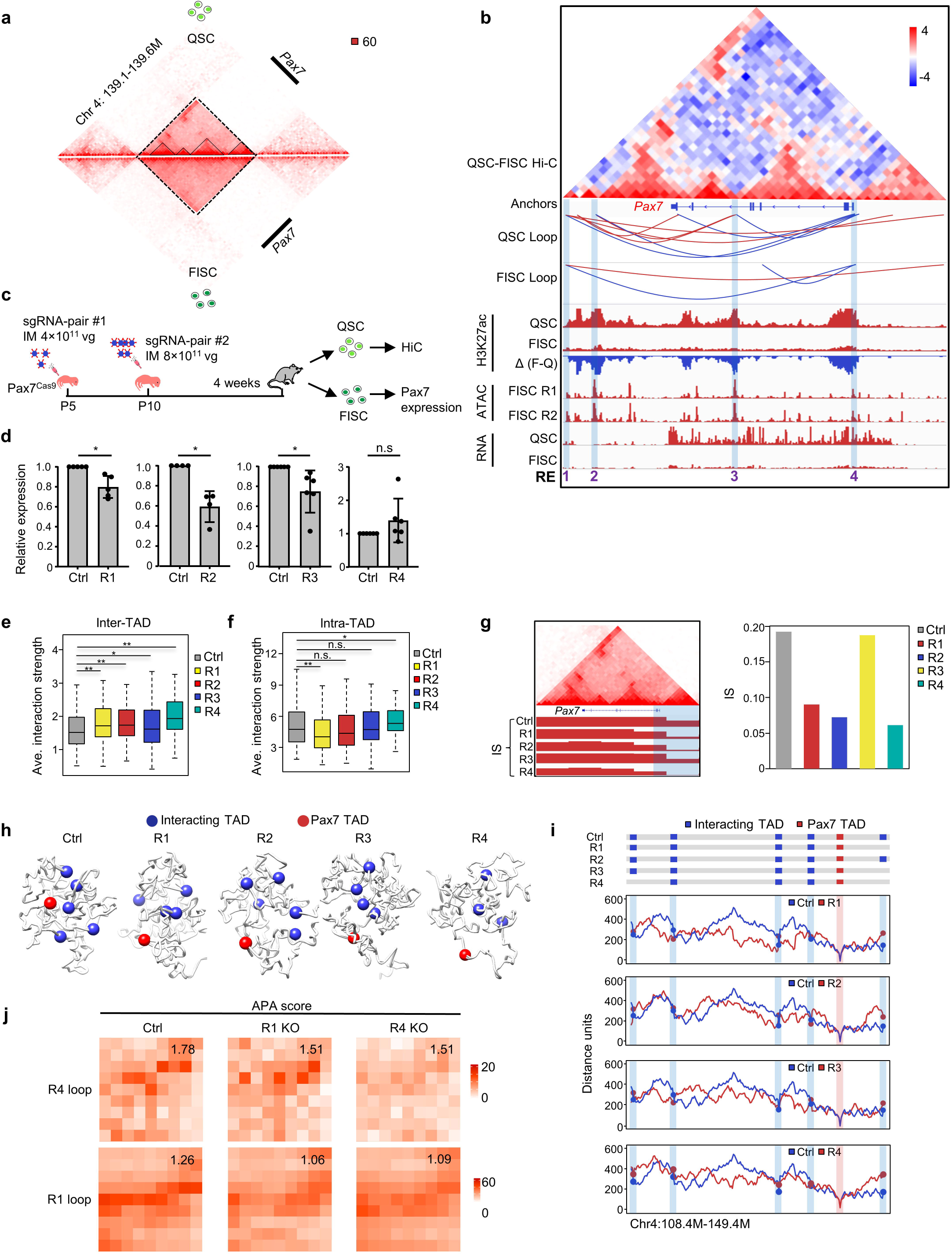
3D regulatory interactions orchestrate Pax7 expression dynamics in SC. **a** Comparison of contact frequencies at the TAD encompassing *Pax7* locus between QSC (Upper) and FISC (Lower). The dashed and solid lines indicate TAD and sub-TADs respectively. **b** Identification of the four regulatory elements (RE, 1, 2, 3 and 4) within the Pax7 residing TAD. Subtraction of FISC from QSC contact matrices shows changes in contact frequencies at the TAD. Chromatin loops identified by FitHiC2, dynamics of RNA expression, and H3K27ac signals surrounding the *Pax7* locus in QSC and FISC, ATAC-seq signal in FISC are shown below. REs are highlighted in green box. **c** Schematic illustration of *in vivo* deletion of each RE in SCs by the CRISPR/Cas9/AAV9-sgRNA system. 4×10^11^gc/mouse of AAV9-sgRNA pair #1 were intramuscularly (IM) injected into Pax7^Cas9^ mice at postnatal day 5 (P5) and followed by 8×10^11^gc/mouse of AAV9-sgRNA pair #2 at P10. The mice were sacrificed for FISC collection and analysis after four weeks. For the control (Ctrl) group, the same dose of AAV9 virus containing pAAV9-sgRNA backbone without any sgRNA insertion was injected. **d** *Pax7* expression was detected by qRT-PCR in the above isolated SCs. The Pax7 expression level was normalized to Hprt1 mRNA level and presented as mean ± s.d. n = 4-6 mice per group. **e-f** *In situ* Hi-C was performed in QSC from Ctrl or each mutant (R1-R4) mouse. Boxplots showing inter- (e) or intra-TAD (f) interactions. **g** (left) Hi-C contact maps (40-kb resolution) of Pax7 TAD in Ctrl(top) and IS (bottom) of Pax7 TADs in Ctrl and sgR1-sgR4. *Pax7* promoter is highlighted in blue box. The TAD boundary covering the *Pax7* promoter is highlighted. (right) Bar plot showing the IS value of the above boundary. **h** 3D chromatin conformation models for the TAD cluster identified with Pax7 residing TAD and other interacting TADs in Ctrl or each mutant QSC. **i** Line plots display distance distribution between *Pax7* residing TAD and interacting TADs. Interacting TADs in Ctrl and sgR1-sgR4 are indicated in blue rectangle (top) and highlighted in blue box (bottom). **j** Aggregated peak analysis (APA) of R4 (top) and R1 (bottom) anchored intra-TAD loops that identified in Ctrl. Numbers on each top right indicate average loop strength. *P<0.05, ***P<0.001. ns, no significance. Data in e and f are presented in boxplots. Center line, median; box limits, upper and lower quartiles; whiskers, 1.5× interquartile range. Statistical analyses were done by T-test and Wilcoxon signed-rank test; *P < 0.01, **P < 0.001, ***P < 0.0001.

To further dissect key regulatory elements governing the above observed 3D re-organization, four putative regulatory elements (REs, R1-R4) were defined based on the looping interactions as well as ATAC-seq, H3K27ac ChIP-seq signals (Fig. 5a). Among them, all four were labeled by high ATAC-seq signals. R4 resided in the *Pax7* promoter region. R2 and R3 were marked with sharp H3K27ac signals in QSC, which disappeared in FISC; R1 resided in the border region. Evident loops between R1/R4, R2/R4, or R3/R4 were observed in QSC but only R1/R4 loop retained in FISC (Fig. 5b). To experimentally validate the functionality of these REs in dedicating Pax7 expression, deletion of each in QSC was achieved by harnessing the muscle specific CRISPR/Cas9/AAV-sgRNA *in vivo* genome editing system recently developed in our group^44^ (Fig. 6c). Briefly, two pairs of sgRNAs targeting the same region (Suppl. Fig. 6a) were applied successively: 4×10^11^ viral genomes (vg)/mouse of AAV9-sgRNA pair #1 was intramuscularly (IM) injected into Pax7^Cas9^ mouse at postnatal age of P5 and followed by 8×10^11^ vg /mouse of AAV9-sgRNA pair #2 at P10. The mice were then sacrificed for SC isolation and analysis after four weeks. For the control (Ctrl) group, the same dose of AAV9 virus containing pAAV9-sgRNA backbone without any sgRNA insertion was injected. The variation in deletion efficiency was detected (Suppl. Fig. 6b): R1 (90%) and R4 (76.1%) manifested more efficient deletion than R2 (15.4%) or R3 (17%), which may be due to the inherent difference of protospacer sequences^45^. As a result, the deletion of R1, R2 or R3 led to 20%, 41%, 25% decrease of Pax7 transcripts in FISC (Fig. 5d). Correspondingly, PAX7 protein levels were also decreased (Suppl. Fig. 6c). Interestingly, removal of R4, the *Pax7* promoter region, did not disrupt Pax7 mRNA level (Fig. 5d) despite ∼ 40% decrease of PAX7 protein (Suppl. Fig. 6c).

**Fig. 6.**
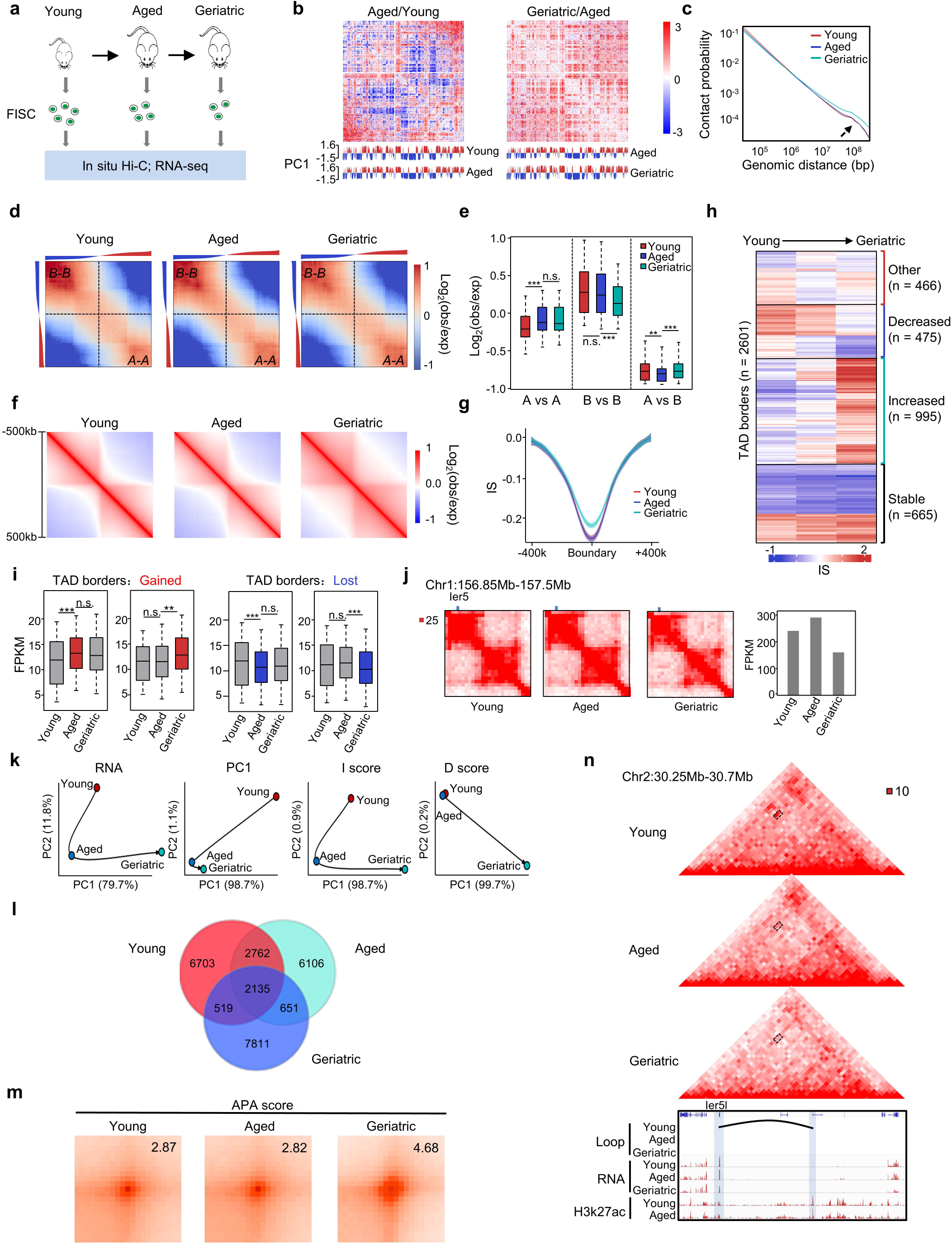
Multiscale analysis of 3D genome during mouse SC aging. **a** Schematic illustration of the aging SC study. FISCs were collected from Young (2 months), Aged (23-24 months) and Geriatric (28-32 months) male Pax7-nGFP mice by FACS sorting. *In situ* Hi-C and RNA-seq were performed in the above cells. **b** The Hi-C data was analyzed and differential heatmaps in Aged/Young (left) or Geriatric/Aged (right) are shown for chromosome 1. The color maps are displayed on the same scale for each comparison. **c** Contact probability relative to genomic distance at each stage. **d** Saddle plots showing the genome-wide compartmentalization over time. Percentage of compartment A/B and PC1 value of each 100-kb loci arranged by eigenvector are shown on top. Corners represent intra-compartment(A-A/B-B) interactions. **e** Contact enrichment of 100-kb loci between compartments of the same (A vs A or B vs B) and different (A vs B) types. P values were calculated by a Wilcoxon signed-rank test. **f** TAD boundaries were identified and aggregate Hi-C maps in a 1Mb region centered on the invariant boundaries during aging. Data are presented as the log ratio of observed and expected contacts in 40kb bins. **g** Average insulation score in an 800 kb region centered on invariant TAD boundaries. Lines denote the mean values, while the shaded ribbons represent the 95% CI. **h** k-means clustering (k = 20) of IS. Bar graphs show IS kinetics for groups that increased (n = 1,146), decreased (n = 528), transiently increased (n = 312) or did not change (n = 674). **i** Boxplots depicting gene expression (RPKM) dynamics for border regions that were gained (left) or lost (right) during SC aging. Stages that gain or lost TAD borders are indicated in blue and right respectively. **j** Illustration of in situ Hi-C contact maps (40-kb resolution) on the *Ier5* locus during SC aging. The bar graphs show the RNA expression of the Ier5. **k** PCA of RNA expression, PC1 values, I-score and D-score during SC aging. Black arrows denote the hypothetical trajectory. **l** Chromatin loops were identified and Venn diagrams showing the overlapping of loops at each stage. **m** APA analysis of chromatin loops detected at young, aged and geriatric FISCs. Bin size, 10 kb. Numbers indicate average loop strengthen. **n** (top) Representative Hi-C contact maps of Chr2: 30.25Mb-30.7Mb depicting the changes in chromatin looping frequencies between *Ier5l* promoter and a putative enhancer during SC aging. Dashed box, chromatin loops between *Ier5l* promoter and putative enhancer. Genome browser tracks showing corresponding RNA-seq, and H3K27ac signals. Light blue box showing the *Ier5l* promoter and a potential enhancer region. Black curve denotes the chromatin loop. Data in e and i are presented in boxplots. Center line, median; box limits, upper and lower quartiles; whiskers, 1.5× interquartile range. Statistical analyses were done by Wilcoxon signed-rank test; *P < 0.01, **P < 0.001, ***P < 0.0001

To further determine the potential effect of the above RE deletion on 3D structure at the *Pax7* locus, *in situ* Hi-C was performed in QSC isolated from the above mice to obtain > 220M uniquely aligned contacts for downstream analysis (Suppl. Fig. 6d and Suppl. Data 7). First, comparison of the Hi-C matrices at 40 kb and 10 kb resolution (Suppl. Fig. 6e) demonstrated that the 3D conformation at *Pax7* locus from the *Pax7*^Cas9^ mice used in this experiment was comparable to that from the earlier used Pax7-nGFP mice (R = 0.85 and 0.83 respectively) (Suppl. Fig. 6f). Next, we found that removal of each RE caused significant increase of the inter-TAD interaction strength genome-wide (Fig. 5e); the intra-TAD interaction strength decreased significantly upon loss of R1 but increased upon R4 loss and unaltered when R2 or R3 was removed (Fig. 5e). Although no obvious effect was observed on TAD organization surrounding *Pax7* (Suppl. Fig. 6g), close examination of the insulation score at the boundary containing *Pax7* promoter revealed a decrease upon loss of R1, R2 or R4 (Fig. 5g and Suppl. Data 7), indicating strengthened border insulation upon removal of these elements. This was not observed when R3 was removed but its deletion led to fusion of nearby sub-TADs into a new sub-TAD (Supp. Fig. 6g). Moreover, a TAD cluster was defined and formed through interaction of the *Pax7* residing TAD with four other TADs in Ctrl QSC; the interaction was disrupted upon loss of each RE (Fig. 5h-5i), which was further supported by virtual-4C using *Pax7* promoter as the viewpoint showing decreased interaction frequency among the TADs (Suppl. Fig. 6h). Interestingly, at the loop level, the interaction strength of R1 and R4 anchored intra-TAD loops were both decreased upon R1 or R4 deletion (Fig. 5j, and Suppl. Data 7). Interestingly, we also observed a decreased number of intra-chromosomal loops anchored in *Pax7* TAD upon deletion of each RE (Suppl. Fig. 6i). Altogether, the above results highlight 3D rewiring at *Pax7* locus during SC activation and unveil key REs dictating the 3D remodeling and Pax7 expression.

### Multiscale analysis of 3D genome during mouse SC aging

It is well accepted that SCs decline in their regenerative capacity with aging^28^, partly ascribed to the intrinsic changes at both transcriptomic and epigenomic levels. However, it is not known whether 3D genome rewiring occurs in SCs during mouse physiological aging. To fill the gap, we collected FISCs from wild-type mice of 2 (young), 23-24 (aged) and 28-32 (geriatric) months of age (Fig. 6a). Consistent with prior findings^27^, a decrease in FISC number was detected in the aged (3.3%) vs. young (4.3%) mice and became more prominent at the geriatric stage (1.8%) (Suppl. Fig. 7a-b). Transcriptional changes were profiled by RNA-seq in the above FISCs with at least two replicates from each stage. In line with previous findings^46^, gene expression profiles were remarkably distinct between young and aged SCs (R = 0.74), and became more divergent in geriatric cells (R = 0.65) (Suppl. Fig. 7c). Notably, *Pax7* expression was steadily upregulated with SC aging, along with stress responsive genes, *Egr1* and *Atf3*; while *Myod* and *Myc*^44^ were highly induced at aged followed by a decline at geriatric phase (Suppl. Fig. 7d). Next, results from pair-wise comparison showed 1,122 genes were significantly up- and 1,016 down-regulated in aged vs young SCs; whereas 910 and 1,104 were in geriatric vs. aged SCs (Suppl. Fig. 7e and Suppl. Data 8). Nevertheless, across the three stages, gene expression adopted distinct dynamics with only a limited proportion of genes continuously increased (n = 120) or decreased (n = 53) (Suppl. Fig. 7f). Consistently, GO analysis revealed distinctly enriched terms in aged and geriatric stages (Supp. Fig. 7g and Suppl. Data 8). For instance, the upregulated genes in aged vs. young SCs were enriched in terms such as “cell adhesion”, “inflammatory response” and “immune response”, while those in geriatric vs. aged SCs were linked with “membrane”, “cell surface” and “extracellular space”.

Next, to delineate the changes in 3D genome organization with SCs aging, *in situ* Hi-C was performed at the above stages with three to four replicates for each to obtain a total of >300 million uniquely aligned contacts; high correlation (R ≥ 0.95) was achieved among the replicates (Suppl. Fig. 8a and Suppl. Data 9). Overall, young SCs exhibited distinct 3D genome organization compared with aged or geriatric SCs; long-range (> 3 Mb) interaction frequencies increased significantly in geriatric SCs compared with young and aged SCs (Fig. 6b-c, Suppl. Fig. 8b). Interestingly, the interaction strength increased between A compartments (A-A) during young to aged transition but remained constant when SCs progressed into geriatric stage; on the contrary, a significant decline in B-B interaction strength was detected during aged to geriatric progression but not young to aged transition (Fig. 6d-e and Suppl. Data 9). Overall, an increase in compartmentalization was observed at aged vs young SCs as revealed by decreased inter-compartment contacts, but then recovered at geriatric stage (Fig. 6d-e). In addition, similar to SC lineage progression (Suppl. Fig. 1i), limited compartment switching was observed: only 6.5% of the genome switched compartment at any two time points with B to A and A to B each accounting for 2.9% and 2.4% respectively (Suppl. Fig. 8c). Overall, the compartmental trajectory showed drastic change at aged vs. young while transcriptomic change occurred late in geriatric stage (Fig. 6k). Whereas the lagged expression change behind the compartment switch was noticed (Suppl. Fig. 8d left), concomitant changes in compartment and gene expression were also identified on some loci (Suppl. Fig. 8d middle and right).

At the TAD level, an average of ∼3,274 – 3,783 TAD borders per stage was identified (Suppl. Fig. 8e). Partitioning of the genome into TADs was largely stable during SC aging as >70% TAD borders (n = 2,660) were detected at all stages (Suppl. Fig. 8e, invariant borders). The aggregate interaction matrix revealed a remarkable loss in insulation strength across the TAD borders in geriatric but not aged stage when compared with young SCs (Fig. 6f), which was supported by IS calculation showing a significant increase at geriatric stage (Fig. 6g, Suppl. Fig. 8f and Suppl. Data 10). Further, quantitative analysis of IS through hierarchical clustering also showed a greater portion (n = 995, 38.3%) of all borders exhibited > 20% increase in IS, while 18.3% (n = 475) showed a loss during SC ageing (Fig. 6h). These data suggested TAD borders largely lost insulation strength when SCs entered into geriatric state. Furthermore, we found that local gene expression underwent significant increase or decrease when TAD borders were gained or lost, respectively (Fig. 6i), which was not observed during SC linage progression (Suppl. Fig. 3f, 3g). Nevertheless, in agreement with the findings in Fig. 2n, D-scores were positively associated with gene expression (Suppl. Fig. 8g); and hierarchical clustering revealed that ∼47% of the TADs (n = 1,754) showed D-score increase or decrease (> 15% difference between endpoints) (Suppl. Fig. 8h and Suppl. Data 10). Additionally, the D-score kinetics was highly correlated with changes in PC1 values (Supp. Fig. 8i) or gene expression (Suppl. Fig. 8j) during the transition between two stages. For instance, the intra-TAD interactions surrounding *Ier5* (Fig. 6j) and *Hox10* (Suppl. Fig. 8k) loci were largely attenuated at geriatric stage, which might result in the concomitant decrease of transcriptional levels from aged to geriatric FISC (Fig. 6j, Suppl. Fig. 8j). Lastly, we performed PCA analysis for RNA expression, compartmentalization, IS and D score. Interestingly, transcription and IS adopted a very similar trajectory showing distinct changes from young to aged to geriatric stages; D-score was largely similar between young and aged stages but underwent a drastic change from aged to geriatric stage; PC1 on the contrary displayed the most prominent change from young to aged stage (Fig. 6k), suggesting genome compartmentalization was altered at aged stage while TAD connectivity changed later at geriatric stage.

Finally, chromatin looping analysis led to identification of a total of 11,180, 10,131 and 11,104 loops in young, aged and geriatric FISCs among which 6,703, 6,106, and 7,811 were stage specific (Fig. 6l and Suppl. Data 10). An increase in interaction strength of chromatin loops was observed when SCs aged (APA score = 4.68 in geriatric vs 2.87 in young and 2.82 in aged) (Fig. 6m). Interestingly, expression changes lagged behind their associated loops, as illustrated on the representative *Ier5l* locus (Fig. 6n). The loop between the promoter and a distal enhancer in young cells disappeared in aged cells but decreased gene expression did not occur until the geriatric stage.

## Discussion

While mechanisms underlying SC lineage progression have been well studied at the transcriptional and epigenetic levels^1, 20, 29, 47^, how genome architecture reorganization contributes to this process remains enigmatic. Here, we examined the genome conformation in SCs at distinct stages and revealed the characteristics of chromatin 3D landscape at different genomic scales and its association with altered gene expression. First, at the large scale, we identified a shift in the ratio between long- and short-range chromatin interactions and remarkable dynamic compartmentalization during SC lineage progression. Loss of long-range contacts at the late stages (D2 and D3) may reflect a heterogeneous nature of SC populations when a portion of cells enter mitosis during which cells are reported to display a rapid fall-off of chromatin contacts at ∼10 Mb^29, 30^; this was not observed at early stages (QSC, FISC and D1) when SCs mostly reside in G0/G1 phases. In addition, SC lineage progression is accompanied with the dynamic changes in compartmentalization strength (Fig. 1d-e), which could be resulted from the alterations in chromatin states. Previous findings have documented the compartment strength increases concurrently with a gain of heterochromatin marks, H3K27me3^18, 48^ or H3K9me3^49^. Indeed, epigenomic profiling in SCs uncovered a significant loss of H3K27ac marks during SC isolation procedure^20^, and a gain of H3K27me3 marks is associated with SC activation^29, 30^. Hence, it will be of great interest to further dissect the connection between the epigenomic changes with the dynamics of genome compartmentalization during SC lineage progression.

Next, at the TAD level, the most striking changes in TAD border strength occur at early activation stage (FISC), which could be a result of the decrease in CTCF expression during the process^20^ to cause the overall detrition in border strength and looping formation as CTCF is known to play key role in chromatin insulation and TAD boundaries^50, 51^. Alternatively, CTCF independent mechanism could contribute to the TAD border reorganization in FISC vs. QSC. For example, previous observations have demonstrated TAD boundaries are also enriched for housekeeping genes, RNA polymerase II, and active chromatin marks^7, 10^ thus active transcription could play a role in shaping the chromatin organization^52^. Nevertheless, gain or loss of borders does not correlate with the overall changes in local gene expression in FISC vs. QSC, arguing transcriptional alterations per se may not be the main contributor for insulation dynamics in this process. Interestingly, the weakened insulation of TAD borders gained strength when SCs enter differentiation, which is reminiscent of findings from neuron development^7^, suggesting TAD borders are dynamically re-organized during cellular differentiation.

Emerging studies indicated interactions among non-contiguous TADs are prevalent^15–17^. In this study, we thus defined TAD clusters as an assembly of ≥ 3 TADs that are fully connected pairwise using a similar approach previously used to annotate TAD cliques^15^. Nevertheless, our defining approach is modified by taking into account of the short-range interactions between TADs. In agreement with previous findings^15^, the long-range inter-TAD interactions (Q3 and Q4) preferentially reside in B compartments and are associated with repressed gene expression but the short-range ones (Q1 and Q2) do not display such preference. Moreover, we found SE-containing TAD clusters are more active in long-range inter-TAD interactions. Concordantly, 3D modeling revealed SE containing TAD clusters reorganize in nuclear space to orchestrate the stage-specific gene expression. These findings complement recent observations that multi-way interactions occur at gene-dense, active, RNA polymerase II-marked regions or regions containing SEs^16, 17^.

Finally, using high-resolution Hi-C datasets to dissect the chromatin looping in FISC and QSC, we revealed the isolation procedure-induced early activation^20^ is accompanied by a great loss of chromatin looping. Reduction of CTCF expression during this process may provide one reason, considering its dominant effect on chromatin looping^50^. Nevertheless, previous study showed that acute attenuation of CTCF in a short time does not lead to drastic changes in gene expression^35^, which contradicts to the expression alterations of a vast spectrum of genes observed in FISC vs. QSC^20^. Hence, CTCF-independent mechanisms may regulate chromatin looping remodeling during the isolation process. Consistent with the overall looping events, the number of enhancer-promoter contacts is largely reduced in FISC vs. QSC, in agreement with the prominent loss in H3K27ac levels during this process^20^. Intriguingly, among the loci we examined, despite a great portion bear enhancer marks in both QSC and FISC, the interaction is restricted to only one stage to enabling the timely gene activation. Our findings, together with previous reports^7, 13, 39, 53^, thus highlight that the interactions provide an additional layer of fine-tuned regulation in lineage development, which cannot be simply inferred from the 1D enhancer profiling. In the future, it will be interesting to identify the factors that drive the decommission of enhancer repertoire and rewiring of E-P contacts during SC early activation.

In addition to the normal SC lineage progression, we delineated the 3D genome remodeling during SC aging. How SCs deteriorate with aging remains largely unknown^54, 55^. Here, we have provided a comprehensive view of how genome 3D configuration in SCs undergo massive changes during the natural aging course of mouse. Interestingly, our finding reveal that despite differences between young and old SCs, geriatric cells display a more prominent gain in long-range contacts and loss of TAD insulation. This is in agreement with recent findings suggesting SCs switch from quiescence to an irreversible pre-senescence state at geriatric age^27^; and a progressive gain of far-cis contacts and loss of insulation has been shown during the onset of various types of cellular senescence^23–25^. However, our results highlighted a preferential gain of A-A interactions and loss of B-B interactions during SC aging, contrasting with previous findings in either oncogene-induced senescence (OIS) or replicative senescence (RS) scenario^25^. In OIS the genome organization is dominated by heterochromatin or B-B interactions resulting in a stronger genome compartmentalization, whereas in RS, dampened A-A interactions causes a decrease in compartmentalization^25^. We speculate these discrepancies may reflect the innate differences between pre-senescence state of geriatric SCs and mature or deep senescent cellular state studied in other works. In addition, SCs need to integrate the information from the deteriorated aging milieu^27, 30, 56^, which may render the state of geriatric SCs more complex and heterogeneous^31^ than the *in vitro* systems used in other studies. Consistent with the notion, PCA analyses revealed distinct trajectories of transcriptome, compartmentalization, TAD insulation and intra-TAD interactions (Fig. 6k). Geriatric SCs overall lose insulation at TAD borders, agreeing with the observations in OIS and RS cells^25^. Recent reports suggest HMGB1 and HMGB2 proteins act as a rheostat of topological insulation upon OIS senescence entry^24, 26^. Consistently, we observed decreased expression of HMGB1 and HMGB2 with SC aging (Suppl. Fig. 7d); it will thus be interestingly to investigate the possible role of HMGB proteins in SC aging. In summary, our study represents the first to describe the 3D genome rewiring in adult stem cells of physiologically aged mammalian organism, and we anticipate more studies from other tissues or organs, will enable us to extract the common paradigm controlling stem cell aging.

The last key contribution of the present study is the elucidation of Pax7 regulation at 3D level. Pax7 represents the uttermost indispensable TF regulating SC maintenance and regenerative capacity^6, 57, 58^. However, current knowledge regarding the upstream regulation of Pax7 expression remains largely unknown. At the transcriptional level, Pax7 RNA expression is vulnerable to diverse environmental cues and rapidly decreases upon SC activation even in response to isolation procedure^20^. Recent studies have provided epigenetic clues that the *Pax7* locus is embedded in a SE region in homeostatic quiescence state and the enhancer signals diminish upon activation^20^, which may account for the acute loss of Pax7 expression. Complementary to these findings, we here uncovered *Pax7* locus is self-organized into a powerful TAD with profound sub-TAD organization identified in quiescent SCs, and the organization largely collapses upon activation. In addition, we identified significant chromatin loops within the Pax7 TAD involving four interconnected REs, including promoter, enhancers, and boundary elements. Leveraging our in-house developed *in vivo* genome editing system, we provide *in vivo* evidence to show that deletion of either of the four REs disturbed Pax7 RNA expression in SCs, and resulted in distinct effects on 3D genome organization at *Pax7* locus. Removal of the downstream boundary region (R1) alone did not cause TAD fusion, possibly due to the enhanced contact newly established between the two convergent CTCF sites^7, 12^. Intriguingly, it instead led to significant reduction of intra-TAD contacts. Deletion of the downstream enhancer region (R2) largely retained the TAD structure with a marginal loss of intra-TAD contacts, especially the interactions between sub-TADs. In contrast, loss of the intronic enhancer region (R3) resulted in more structured sub-TAD patterns. In contrast to the above deletions resulting in a consistent decreased Pax7 expression caused by the above deletions, deletion of the promoter region (R4), surprisingly, enhanced Pax7 RNA expression, which may be caused by the use of an alternative TSS downstream of the enhancer region. Moreover, we showed that the Pax7 TAD is also involved in inter-TAD interactions to form a TAD cluster. Deletion of each RE within the TAD also abrogated the clustering. Altogether, our findings for the first time unveil that Pax7 transcription is tightly controlled by the embedded REs and 3D interactions. Nevertheless, we need to point out that the editing efficiency on each RE varied from 15.4% for R2 to 90% for R1, which may be intrinsically dictated by the *in vivo* editing system. The It is possible that more evident changes of chromatin architecture and Pax7 expression can be achieved if the editing efficiency reaches 100% equally for all REs using traditional knockout models.

In conclusion, our study provides a comprehensive view of 3D genome organization in SC lineage progression and SC aging. Our findings demonstrate that chromatin rewiring at different genomic scales underpin the transcriptome remodeling and SC activities, underscoring the importance of 3D regulation in stem cell lineage development and stem cell aging. The datasets produced in this study for the first time provide a rich resource of chromatin interactions for the SC community. In addition, our work uncovers the 3D regulation of PAX7, underscoring the importance to connect the 3D genome rewiring to the fine-tuned TF dynamics during lineage development.

## Methods

### Mice

All animal experiments were performed following the guidelines for experimentation with laboratory animals set in the Chinese University of Hong Kong and approved by the Animal Experimentation Ethics Committee (AEEC) of the Chinese University of Hong Kong (CUHK). The mice were maintained in animal room with 12 hrs light/12 hrs dark cycles at Animal Facility in CUHK. For SC lineage development study, two-month-old male Pax7-nGFP mice were used to isolate satellite cells. For aging SC study, male wild-type C57BL/6 mice of 2 (young), 23-24 (aged) and 29-31 (geriatric) months of age were used to isolate SCs. For *Pax7* locus dissection, Pax7^Cas9^ mice were generated by crossing homozygous Cre-dependent Rosa26^Cas9-EGFP^ knockin mouse (B6;129-Gt (ROSA)26Sor^tm1(CAG-cas9*-EGFP) Fezh/J^; stock number 024857, Jackson Laboratory) with a Pax7^Cre^ strain, as previously described^44^.

### Satellite cell isolation by FACS

SCs were sorted by FACS based on established methods^59^. Briefly, hindlimb muscles from Pax7-nGFP mice were collected and digested with collagenase II (1000 U ml^-1^, Worthington) for 90 min at 37°C, and then the digested muscles were triturated and washed in washing medium (Hams F-10 media (Sigma), 10% HIHS (Gibco), penicillin/streptomycin (1×, Gibco) before SCs were liberated by treating with Collagenase II (1000 U ml^-1^) and Dispase (11 U ml^-1^) for 30 min at 37°C. Mononuclear cells were filtered with a 40-µm cell strainer and GFP+ SCs were sorted out by BD FACSAria Fusion cell sorter (BD Biosciences). For SC isolation from C57BL/6 mice, the filtered mononuclear cells were additionally incubated with Vcam1-biotin (105704, BioLegend), CD31-FITC (102506, BioLegend), CD45-FITC (103108, BioLegend), and Sca1-Alxa647 (108118, BioLegend). The Vcam1 signal was amplified with Streptavidin-PE (554061, BD Biosciences). All antibodies were used at a dilution of 1:75. Vcam1+, CD31-, CD45-, Sca1- cells were sorted out by BD FACSAria Fusion cell sorter (BD Biosciences). For QSC isolation, we followed the procedures described previously^20^. Briefly, the muscles were pre-fixed in ice-cold 0.5% paraformaldehyde (PFA) before cell dissociation. Collagenase II and Dispase were used at 2,000 U mL^-1^ and 22 U ml^-1^ for the digestion respectively.

### AAV9 virus production, purification and injection

AAV9 virus was produced as described previously^44, 60^. In brief, HEK293FT cells were seeded in T75 flask and transiently transfected with AAV9-sgRNA vector (5 μg), AAV9 serotype plasmid (5 μg), and pDF6 (AAV helper plasmid) (10 μg) at a ratio of 1:1:2 using polyethyleneimine (PEI) when the cell reached 80%∼90% confluency. Twenty-four hours after transfection, the cells were changed to growth medium (DMEM with 10% FBS, 100 U ml^-1^ penicillin, 100 μg ml^-1^ streptomycin and 2 mM L-glutamine) and cultured for another 48 hrs. The cells were harvested and washed with PBS two times. To release the AAV9 virus, the pellet was re-suspended with lysis buffer (50 mM Tris-HCl pH 8.0, 150 mM NaCl) followed by three sequential freeze–thaw cycles (liquid nitrogen/37°C). The lysates were treated with Benzonase (Sigma) together with MgCl_2_ (final concentration: 1.6 mM) at 37°C for 1 hr followed by centrifugation at 3,000 rpm for 10 min. The supernatant was filtered with 0.45 μm sterile filter and added with equal volume of 1 M NaCl and 20% PEG8000 (w/v) to precipitate the virus at 4 °C overnight. After centrifugation at 12,000 g for 30 min at 4 °C, the pellet was re-suspended with sterile PBS and then subject to centrifugation at 3,000 g for 10 min. Equal volume of chloroform was then added and shaken followed by spin down at 12,000 g for 15 min at 4 °C. The aqueous layer was filtered by 0.22 μm sterile filter and passed through a 100 kDa MWCO (Millipore). The concentrated solution was washed with sterile PBS three times. The tilter of the AAV9 virus was determined by qRT-PCR using primers targeting the CMV promoter. Sequences of the used primers are listed in Suppl. Table 1. Heterozygous Pax7^Cas9^ mice were intramuscularly (IM) injected with 4×10^11^ viral genomes (vg) of AAV9-sgRNA pair #1 at postnatal day 5 (P5) and followed by 8×10^11^ vg of AAV9-sgRNA pair #2 at P10. SCs were isolated 4 weeks post AAV injection.

### Cell culture

HEK293FT cells were obtained from ATCC (CRL-3216) and maintained in DMEM supplemented with 10% FBS, 100 U ml^-1^ penicillin and 100 μg of streptomycin in a 5% CO_2_ humidified incubator at 37°C. Freshly-isolated SCs were cultured in Hams F10 medium (Sigma) with 20% FBS, penicillin/streptomycin (1×) and 2.5 ng ml^-1^ basic fibroblast growth factor (bFGF, 13256, Life Technologies).

### Plasmids

Site-specific sgRNAs were selected using a web tool Crispor^61^ (http://crispor.tefor.net/). Synthesized oligonucleotides encoding guide sequences were cloned into the AAV9-sgRNA transfer vector (AAV: ITR-U6-sgRNA(backbone)-CMV-DsRed-WPRE-hGHpA-ITR) using Sap I site. To generate dual AAV-sgRNAs expression plasmid, the second sgRNA was constructed into the AAV9-sgRNA vector together with the gRNA cassette and U6 promoter using Xba I and Kpn I sites. Sequences for sgRNAs and primers used to detect deletion efficiency are listed in Suppl. Table 1.

### RNA isolation, quantitative RT-PCR and RNA-seq

For FISC and cultured SCs, total RNAs were extracted using TRIzol reagent (Invitogen). For QSC, total RNAs were isolated from the above-described pre-fixed cells using a miRNeasy FFPE Kit (Qiagen) according to the manufacturer’s instruction. For quantitative RT-PCR, cDNAs were prepared using HiScript III 1^st^ Strand cDNA Synthesis Kit (Vazyme, R312-01). Real-time PCR reactions were performed on a LightCycler® 480 Instrument II (Roche Life Science) using Luna Universal qPCR Master Mix (NEB, M3003L). Sequences of all primers used are listed in Suppl. Table 1. For polyA+ mRNA-seq, total RNAs were subject to poly(A) selection (Ambion, 61006) followed by library preparation using NEBNext® Ultra™ II RNA Library Preparation Kit (NEB, E7770S). Libraries were paired-end sequenced with read lengths of 150 bp on an Illumina HiSeq X Ten instrument. For total RNA-seq, total RNAs were isolated from QSC, FISC or cultured SCs; library preparation and paired-end sequencing was performed by Beijing Genomics Institute (BGI, Hong Kong) with read length of 100 bp on Nextseq 500.

### Immunoblotting and immunofluorescence

These were performed according to our standard procedures^62–65^. Briefly, total proteins were extracted using RIPA lysis buffer. The following dilutions of antibodies were used for each antibody: PAX7 (Developmental Studies Hybridoma Bank; 1:1000), GAPDH (Santa Cruz Biotechnology, sc-137179, 1:2000). For immunofluorescence staining, following antibodies and related dilutions were used: PAX7 (Developmental Studies Hybridoma Bank; 1:100). All fluorescent images were captured with a fluorescence microscope (Leica DM6000 B).

### Chromatin immunoprecipitation followed by high-throughput sequencing (ChIP-seq)

Following our standard procedure^44, 62, 63, 66^, approximately 300,000-600,000 FISCs were cross-linked with 1% formaldehyde for 10 minutes at room temperature and quenched by 125 mM Glycine. Cells were washed and collected by centrifugation at 700 g for 5 min at 4°C, flash-frozen in liquid nitrogen and stored at – 80°C. Cells were lysed in 1 ml of LB1 (50 mM HEPES-KOH pH 7.5, 140 mM NaCl, 1 mM EDTA, 0.25% Triton X-100, 0.5% NP-40, 10% glycerol, supplemented with cOmplete protease inhibitors (Roche)) and incubated at 4°C for 10 min. Following centrifugation at 1,350 g for 5 min at 4°C, the pellets were washed with 1 ml of LB2 (10 mM Tris-HCl pH 8.0, 200 mM NaCl, 1 mM EDTA pH 8.0, 1 mM EGTA pH 8.0, supplemented with complete proteinase inhibitors) by incubating at 4°C for 10 min. Following centrifugation at 1,350 g for 5 min at 4°C, nuclei were rinsed twice with 1 ml of sonication buffer (10 mM Tris-HCl pH 8.0, 0.1% SDS, 1 mM EDTA, supplemented with complete proteinase inhibitors). Then, the nuclei were re-suspended in sonication buffer and sonicated with Covaris S220 (Intensity 140W, Duty Cycle 5%, Cycles per Burst 200, Time 7 mins). The resulting lysate was supplied with NaCl and Triton X 100 to reach a final concentration at 150 mM NaCl and 1% Triton X 100 and then cleared by centrifugation for 20 min at 20,000 g and then incubated with washed Dynabeads Protein G (Invitrogen, 10004D) for 2 hrs at 4°C. The cleared lysate was incubated with 0.75 μg H3K27ac (ab4729, Abcam) or IgG control (sc-2027, Santa Cruz) for overnight at 4°C. On the next day, 10 μl Dynabeads Protein G was washed with IP buffer (10 mM Tris-HCl pH 8.0, 150 mM NaCl, 0.1% SDS, 1 mM EDTA, 1% Triton X100, supplemented with complete proteinase inhibitors) and then applied to each IP reaction followed by incubation at 4°C for another 2 hrs. Beads were then collected and washed twice with IP buffer, two times with High salt wash buffer (10 mM Tris-HCl pH 8.0, 500 mM NaCl, 0.1% SDS, 1 mM EDTA, 1% Triton X100, supplemented with complete proteinase inhibitors), two times with LiCl wash buffer (10 mM Tris-HCl pH 8, 250 mM LiCl, 1 mM EDTA, 0.5% NP-40, 0.5% deoxycholate, supplemented with complete proteinase inhibitors) and one time with cold TE buffer (10 mM Tris-HCl pH8.0, 1 mM EDTA, 50 mM NaCl). Immunocomplexes were eluted in ChIP elution buffer (50 mM Tris-HCl pH8.0, 10 mM EDTA, 1% SDS) with incubation at 65°C for 30 min and eluents were reverse crosslinked by incubating at 65°C for 16 hrs. Immunoprecipitated DNA was treated with RNase A (0.2 mg/ml) at 37°C for 2 hrs, followed by Proteinase K treatment (0.2 mg ml^-1^) at 55°C for 3 hrs. Immunoprecipitated DNA was then subject to phenol:chloroform extraction and ethanol precipitation and finally resuspended in 1x TE buffer. Libraries were prepared using NEBNext® Ultra™ II DNA Library Preparation Kit (NEB, E7645S) and paired-end sequenced with read lengths of 150 bp on an Illumina HiSeq X Ten instrument.

### *In situ* Hi-C library preparation and sequencing

Generation of Hi-C libraries with a low cell number of SCs was performed according to modified protocols^8^. Approximately 100,000 SCs were cross-linked with 1% formaldehyde for 10 minutes at room temperature and quenched by 0.2 M Glycine. The samples were lysed in 500 μL of ice-cold Hi-C lysis buffer (10 mM Tris-HCl pH 8.0, 10 mM NaCl, 0.2% Igepal CA630, supplemented with complete proteinase inhibitors) on ice for 15 min. Samples were then centrifuged at 2,500 g for 5 min. Pelleted nuclei were washed once with 500 μL of 1.25 × NEBuffer 3.1. The supernatant was discarded and the nuclei was re-suspended in 358 μL of 1.25 × NEBuffer 3.1 and add 11 μL of 10% SDS followed by incubation at 37°C for 1 hr. After incubating, 75 μL of 10% Triton X-100 was added to quench the SDS and then incubated at 37°C for 1 hr. 100 U of DpnII restriction enzyme (NEB, R0543) was added and chromatins were digested at 37°C for overnight. Samples were incubated at 62°C for 20 min to inactivate DpnII and then cooled to room temperature. Samples were then centrifuged at 2,500 g for 5 min. Pelleted nuclei were re-suspended in 50 μL of fill-in master mix (3.75 μL of 0.4 mM biotin-14-dATP, 1.5 μL of 1 mM dCTP, 1.5 μL of 1 mM dGTP, 1.5 μL of 1 mM dTTP mix, 2 μL of 5 U μL^-1^ DNA polymerase I, Large (Klenow) Fragment, 1 × NEBuffer 3.1) to fill in the restriction fragment overhangs and mark the DNA ends with biotin. Samples were mixed by pipetting and incubated at 23°C for 4 hrs. Ligation master mix (398 μL of water, 50 μL of 10 × NEB T4 DNA ligase buffer, 1 μL of 50 mg/ml bovine serum albumin, 1 μL of 400 U μL^-1^ T4 DNA ligase) was added and samples were incubated at 16°C for overnight in ThermoMixer C with interval shake. Nuclei were pelleted by centrifugation at 2,500 g for 5 min and 380 μL of the supernatant was discarded. Pellets were then re-suspended in the remaining 120 μL ligation mix supplemented with 12 μL of 10% SDS and 5 μL of 20 mg ml^-1^ proteinase K and incubated at 55°C for 2 hrs with shaking at 1,000 rpm. 15 μL of 5M NaCl was added and the reaction was incubated at 65°C for 16 hrs. DNA samples were subjected to phenol: chloroform extraction and ethanol precipitation and finally re-suspended in 130 μL 0.1 × TE buffer. The purified DNA samples were then sheared to a length of ∼300 bp using Covaris S220 instrument (Intensity 175W, Duty Cycle 10%, Cycles per Burst 200, Time 150 s). The biotinylated DNA was pulled down by 10 μL of 10 mg/ml Dynabeads MyOne Streptavidin C1 beads (Life technologies, 65001). The beads were then re-suspended in 23 μL of 10 mM Tris-Cl pH 8.0 and libraries were prepared by on-bead reactions using NEBNext® Ultra™ II DNA Library Preparation Kit (NEB, E7645S). The beads were separated on a magnetic stand and the supernatant was discarded. After washes, the beads were re-suspended in 20 μL of 10 mM Tris buffer and boiled at 98°C for 10 min. The elute was amplified for 10-13 cycles of PCR with Phanta Master Mix (Vazyme, P511-01) and the PCR products were purified using VAHTS DNA Clean Beads (Vazyme, N411-01). The Hi-C libraries were paired-end sequenced with read lengths of 150 bp on an Illumina HiSeq X Ten instrument.

### Gene expression analysis of RNA-seq data

The raw reads of total RNA-seq were processed following the procedures described in our previous publication^62, 63, 67^. Briefly, the adapter and low-quality sequences were trimmed from 3’ to 5’ ends for each read and the reads shorter than 50 bp were discarded. The reads that passed the quality control were mapped to mouse genome (mm9) with TopHat2. Cufflinks were then used to estimate transcript abundance in Fragments Per Kilobase per Million (FPKM). Genes were annotated as differentially expressed if the change of expression level is greater than 2 folds between two stages/conditions.

### ChIP-seq data analysis

Raw ChIP-seq reads were processed as previously described^66^. Briefly, the adapter and low-quality sequences were trimmed from 3’ to 5’ ends by Trimmomatic and the reads shorter than 36 bp were discarded. Subsequently, the preprocessed reads were aligned to the mouse genome (mm9) using Bowtie2. The aligned reads were then converted to bam format using samtools, and the duplicate reads were removed by Picard (http://broadinstitute.github.io/picard). Peaks were then identified by MACS2 with q-value equal to 0.01 by using the IgG control sample as background. H3K27ac CHIP-seq is used for typical enhancer (TE) and Super enhancers (SEs) identification. For typical enhancer (TE) identification, blacklisted regions from ENCODE were first excluded, and the peaks overlapping with +/- 2kb regions centered at Transcription Start Site (TSS) from Refseq (June 2015) were also discarded. The filtered peaks were defined as TEs and assigned to the expressed transcripts (RPKM >0.5) whose TSS are nearest to the center of the enhancer regions. Super enhancers (SEs) were identified as previously described with minor adjustment^68^. Only the enhancers within 12.5 kb of each another and can be assigned to the same genes were stitched together, then subject to the ROSE algorithm for SE identification.

### *In situ* Hi-C data processing

The *in-situ* Hi-C data was processed with a standard pipeline HiC-Pro^33^. First, adaptor sequences and poor-quality reads were removed using Trimmomatic (ILLUMINACLIP: TruSeq3-PE-2.fa:2:30:10; SLIDINGWINDOW: 4:15; MINLEN:50). The filtered reads were then aligned to reference genome (mm9) in two steps: 1) global alignment was first conducted for all pair-end reads, 2) the unaligned reads were split into prospective fragments using restriction enzyme recognition sequence (GATCGATC) and aligned again. All aligned reads were then merged together and assigned to restriction fragment, while low quality (MAPQ<30) or multiple alignment reads were discarded. Invalid fragments including unpaired fragments (singleton), juxtaposed fragments (re-legation pairs), un-ligated fragments (dangling end), self-circularized fragments (self-cycle), and PCR duplicates were removed from each biological replicate. The remaining validate pairs from all replicates of each stage were then merged, followed by read depth normalization using HOMER and matrix balancing using iterative correction and eigenvector decomposition (ICE) normalization to obtain comparable interaction matrix between different stages.

### Identification and analysis of compartments and TADs

Following previous procedure^8^, **t**o separate the genome into A and B compartments, the ICE normalized intra-chromosomal interaction matrices at 100-kb resolution were transformed to observe/expect contact matrices, and the background (expected) contact matrices were generated to eliminate bias caused by distance-dependent decay of interaction frequency and read depth difference. Pearson correlation was then applied to the transformed matrices and the first principal component (PC1) of these matrices was divided into two clusters. The annotation of genes and the expression profile were used to assign positive PC1 value to gene-rich component as compartment A and negative PC1 value to gene-poor component as compartment B. The compartmentalization strength was measured using cooltools (https://github.com/mirnylab/cooltools). Briefly, all 100-kb compartment bins were divided into 50 equal degrees based on the ranking of PC1 values, and the average interaction strength (observed/expected) was calculated between pairs of 100-kb loci arranged by their PC1 and normalized by genomic distance. The dynamics of interaction strength for intra/inter compartment (A vs. A, B vs. B, A vs. B) were measured using top 20% of compartment bins from both compartments A and B based on absolute PC1 value. Normalized contact matrix at 40 kb resolution of each time point was used for TAD identification using TopDom^37^. In brief, for each 40-kb bin across the genome, a signal of the average interaction frequency of all pairs of genome regions within a distinct window centered on this bin was calculated, thus TAD boundary was identified with local minimal signal within certain window. The false detected TADs without local interaction aggregation were filtered out by statistical testing. Invariant TADs were defined using following criteria: 1) the distance of both TAD boundaries between two conditions is no more than 40 kb; 2) the overlapping between two TADs should be larger than 80%; stage-specific TADs were defined otherwise. The insulation index for each bin was generated based on a previously described method^69^. In brief, the insulation index for each bin was obtained by calculating the number of interaction across a specific bin within certain distance. The insulation score of the identified TAD border was also defined as previously described^69^, which used the local maximum on the outside of TAD to minus the local minimum on the inside of TAD of each boundary bin. The domain score was calculated for each TAD at 40 kb interaction matrix through dividing total intra-TAD contacts by all contacts involving the TAD.

### Identification of TAD-TAD interactions and TAD clusters

The significant TAD-TAD interactions and TAD clusters were identified as previously described^70^. First, the significance of TAD-TAD interactions was tested by applying a non-central hypergeometric (NCHG) model for all pairs of TADs, then corrected by multiple testing with false discovery rate (FDR) < 1% using Benjamini-Hochberg method. Significant TAD-TAD interactions were defined for those TAD pairs with adjusted p-value <0.05 and five-fold enrichment of observed interactions over expected. All significant TAD pairs were then used to identify TAD clusters using the NetworkX Python library (http://networkx.github.io/), where each TAD was represented by a node and interactions were represented by edges. Maximal size of TAD clusters was calculated using the Bron-Kerbosch algorithm and allvial package (https://github.com/mbojan/alluvial) was used to plot the TAD cluster dynamics.

### 3D modeling of TAD cluster

The normalized interaction matrix containing TAD cluster as well as their genomic 3D context were used to build 3D modelling following previous method with modification^71^. (1) to select key elements contained in the region (that is, TADs in selected TAD cluster); (2) to retrieve the top 1% interactors of each of these elements; (3) to create a network between interactors where edges correspond to top 1% interactions between any of the selected interactions; (4) to group the networks allowing the groups in which the ratio (number of edges)/(number of nodes) smaller than 5 were filtered; and (5) to extract the connected groups that contain the most key elements. Next, the normalized interaction matrices of selected regions were modeled using TADdyn^72^. A similar protocol as previously described was used^14^. Briefly, a total of 1000 models were generated for each genomic region and dataset. Each ensemble of models was next clustered on the basis of structural similarity. The absence of major structural differences between clusters prompted us to use all of them for further analysis. Next, TADbit^73^ was used to measure the following features of the models: (1) distance between particles containing genomic regions of interest in the model ensemble; (2) distance distribution between selected pairs or particles; and (3) notable differential distance distributions assessed by the two-sample Kolmogorov–Smirnov statistic. Finally, model images were generated with Chimera^74^.

### Identification of chromatin loops

The loops were identified by FitHIC2^42^ (https://ay-lab.github.io/fithic/) at 5 kb resolution of interaction matrix with p-value<1e-5 and required a five-fold enrichment of observed contacts over expected based on genomic distance. To obtain stage-specific loops, fold change (FC) of interaction strength (observed/expected) between two stages was calculated for each identified loop. Then the top 30% dynamics loops with absolute FC > 2 were defined as stage-specific loops.

### Motif scanning

The position-specific probability matrix of motifs was obtained from the TRANSFAC database^75^. Utilizing FIMO, the TAD boundaries as well as enhancers and promoter regions (+/- 2kb) of PAX7 were scanned for motifs of all expressed transcription factors in both QSC and FISC. The motif was kept if the p-value was smaller than 10^−4^.

### PCA analysis

Principal component analysis (PCA) was generated using the prcomp package in R (V 3.6.2) with FPKM for RNA-seq, PC1 for A/B compartments and domain score/insulation score for conserved TADs. The eigenvalue for the first two components were used, and the explanatory ratio of each component was calculated using the eigenvalue divided by total eigenvalue.

## Data availability

*In situ* Hi-C and RNA-seq data reported in this paper are deposited in the Gene Expression Omnibus database under accession GSE189842. All used datasets from other publications are summarized in Suppl. Table 2. All other data supporting the findings of this study are available from the corresponding author on reasonable request. The source data for Figs 1h, i, 2b-e, k, l, 3b, c, 4a, 5g, j and 6f-h, k-m Suppl. Figs 2a, c, 3a, b 5a, b, f, 7f, g, 8f-h are provided in the Source Data file.

## Acknowledgments

We thank Prof. Zhenyu Ju (Jinan University, China) for providing the aged mice and Prof. Danny Leung (HKUST, Hong Kong, China) for assistance on the Hi-C sequencing. This work was supported by General Research Funds (GRF) from the Research Grants Council (RGC) of the Hong Kong Special Administrative Region (14116918, 14120420, and 14120619 to H.S.; 14115319, 14100018, 14100620, 14106117 and 14106521 to H.W.); the National Natural Science Foundation of China (NSFC) to H.W. (Project code: 31871304); Collaborative Research Fund (CRF) from RGC to H.W. (C6018-19GF); NSFC/RGC Joint Research Scheme to H.S. (Project code: N_CUHK 413/18); Hong Kong Epigenomics Project (EpiHK) Fund to H.W. and H.S.; Area of Excellence Scheme (AoE) from RGC (Project number: AoE/M-402/20).

## Author Contributions

H.W., Y.Z., Y.D. and H.S. designed the project; Y.Z., L.H., Y.L., and X.C. performed the research; Y.D. performed bioinformatic analyses of sequencing data; Y.Z., Y.D., L.H. and H.W. wrote the paper. All authors read and approved the final manuscript.

---

**Suppl. Fig. 1.**
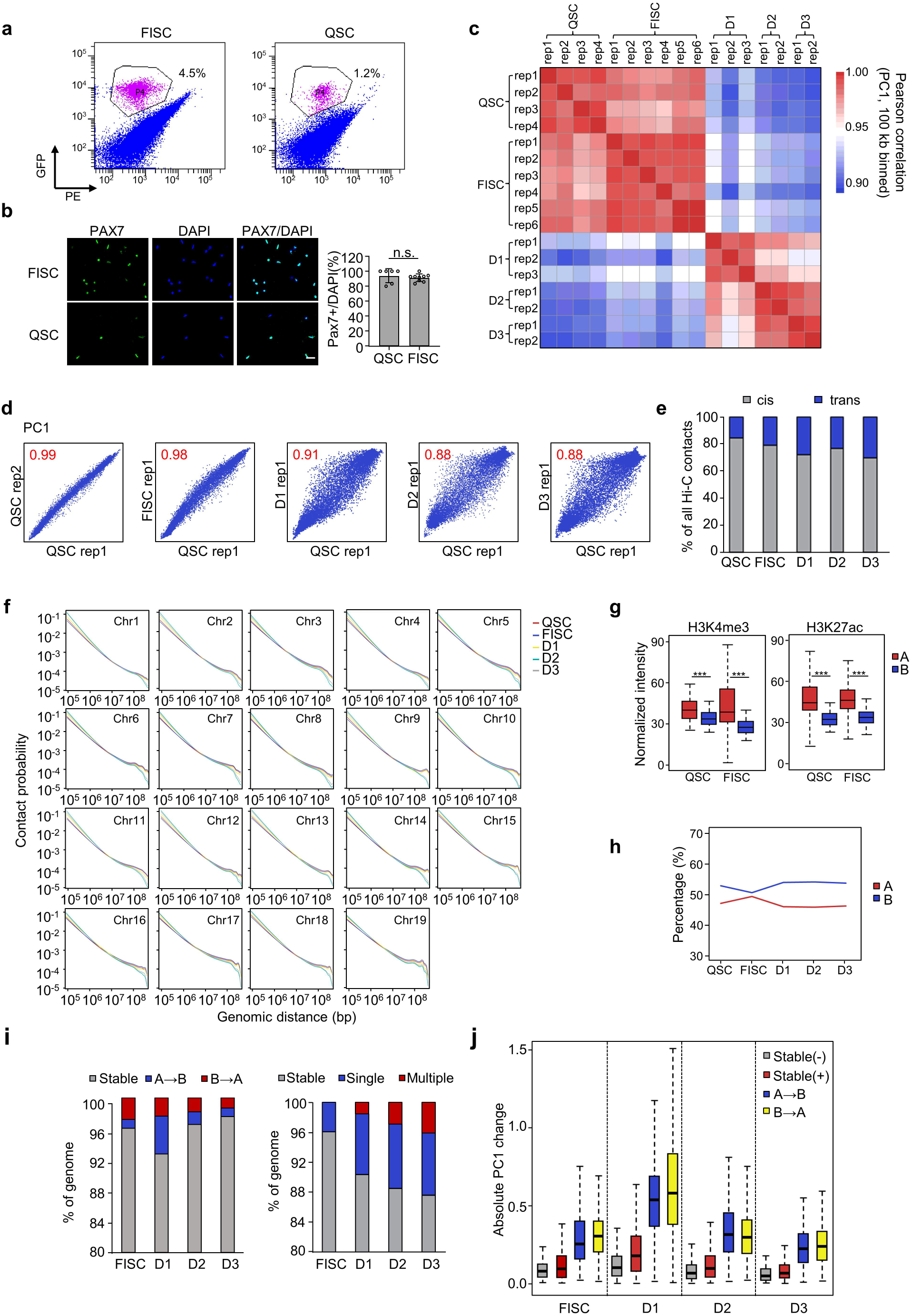

**Suppl. Fig. 2.**
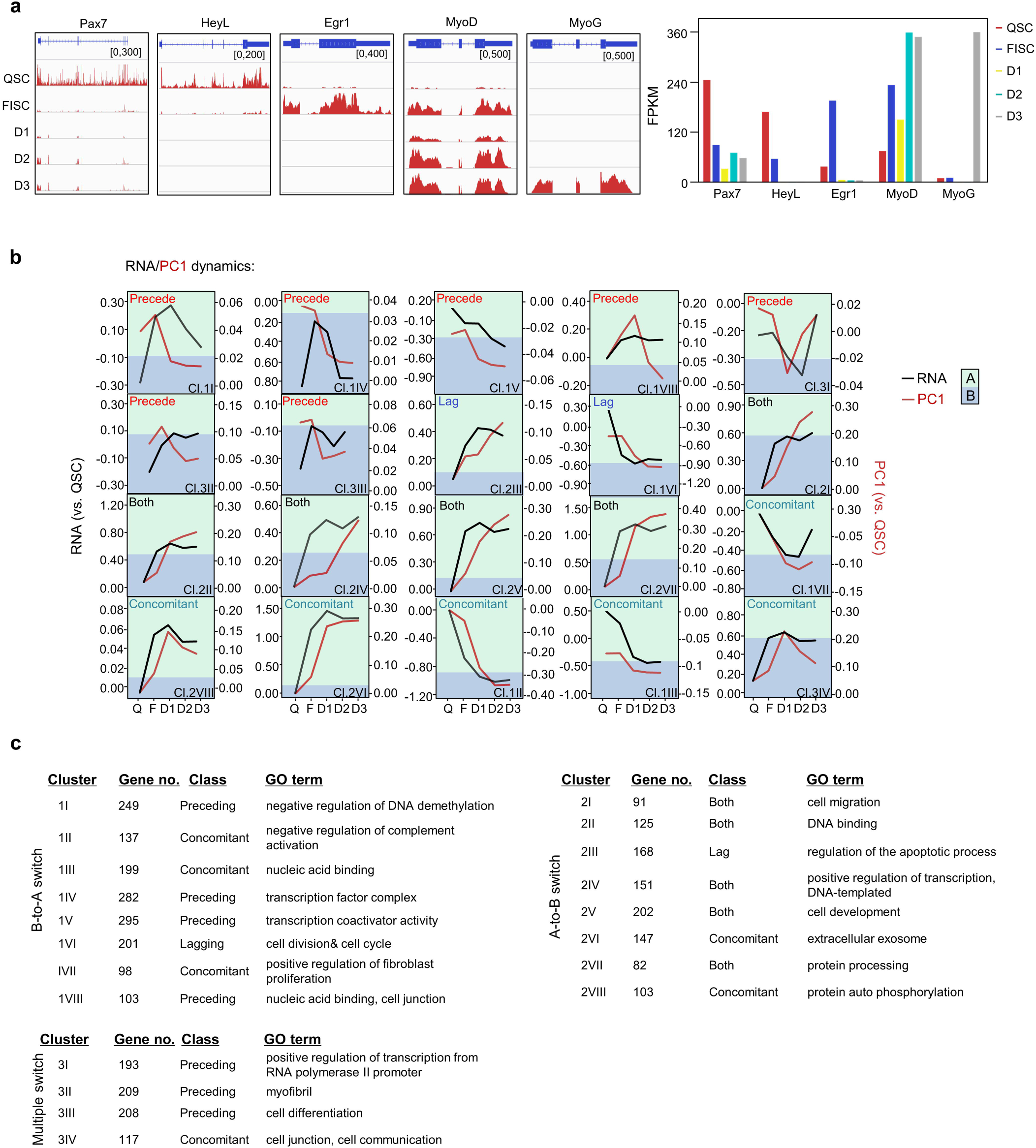

**Suppl. Fig. 3.**
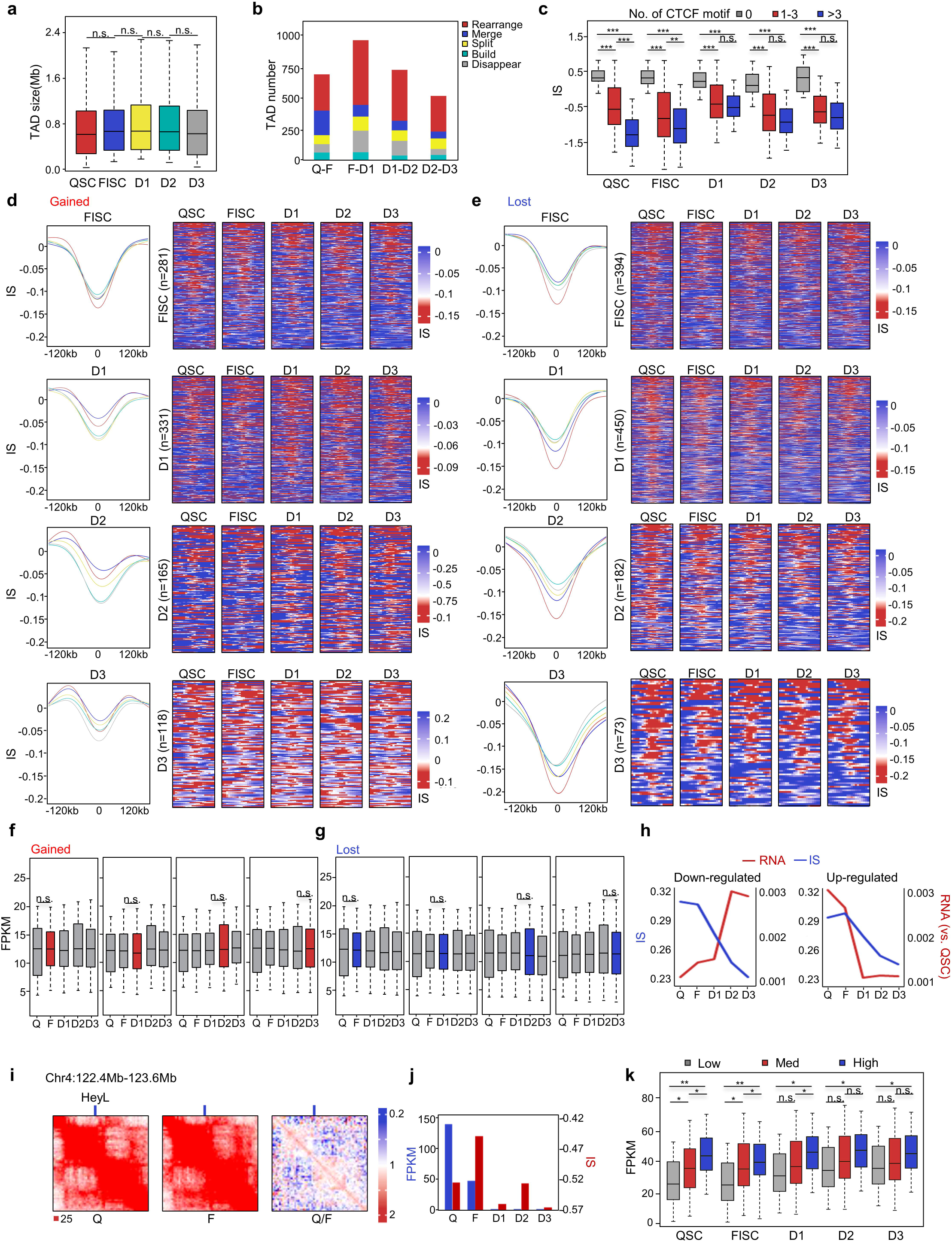

**Suppl. Fig. 4.**
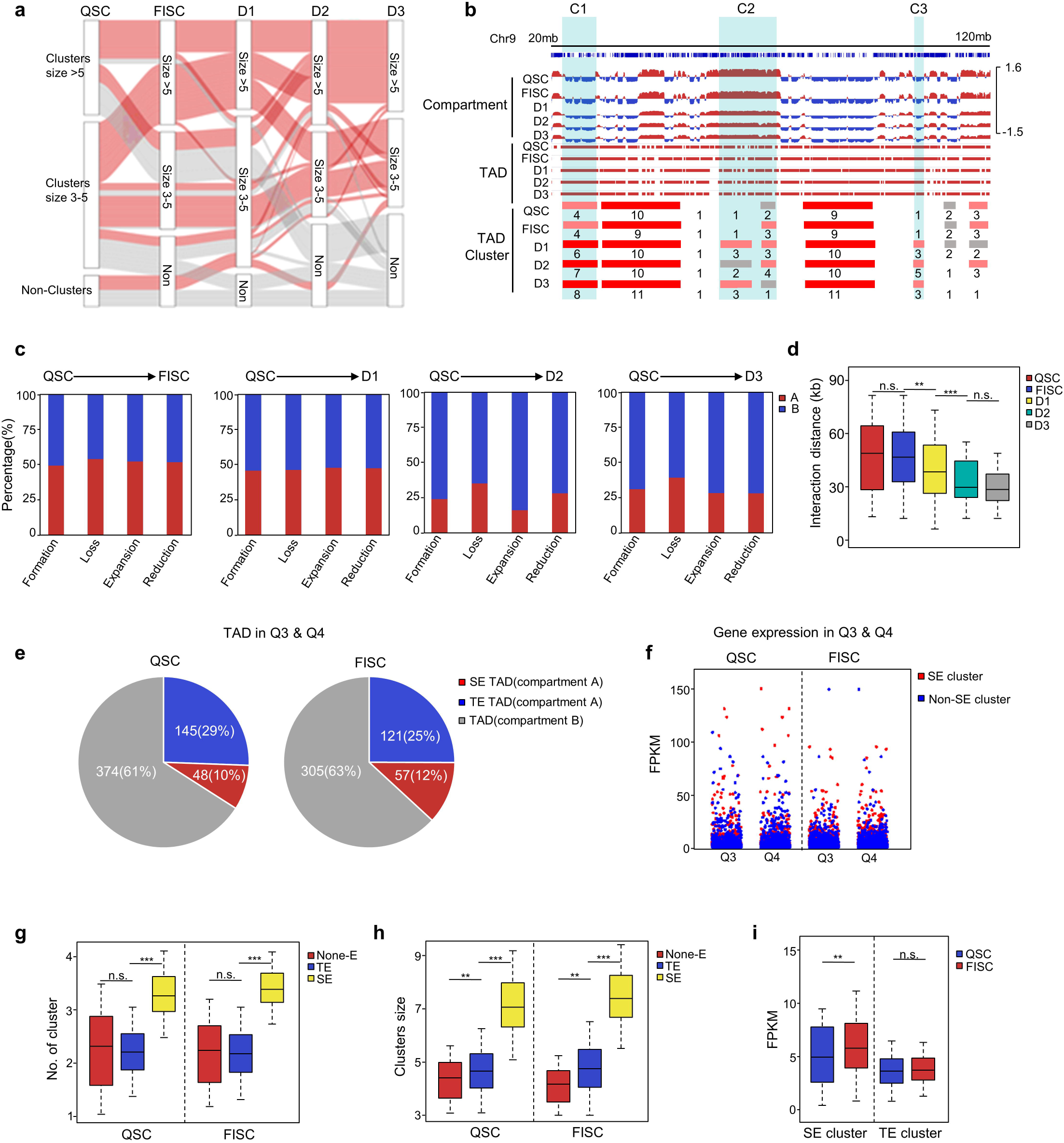

**Suppl Fig. 5.**
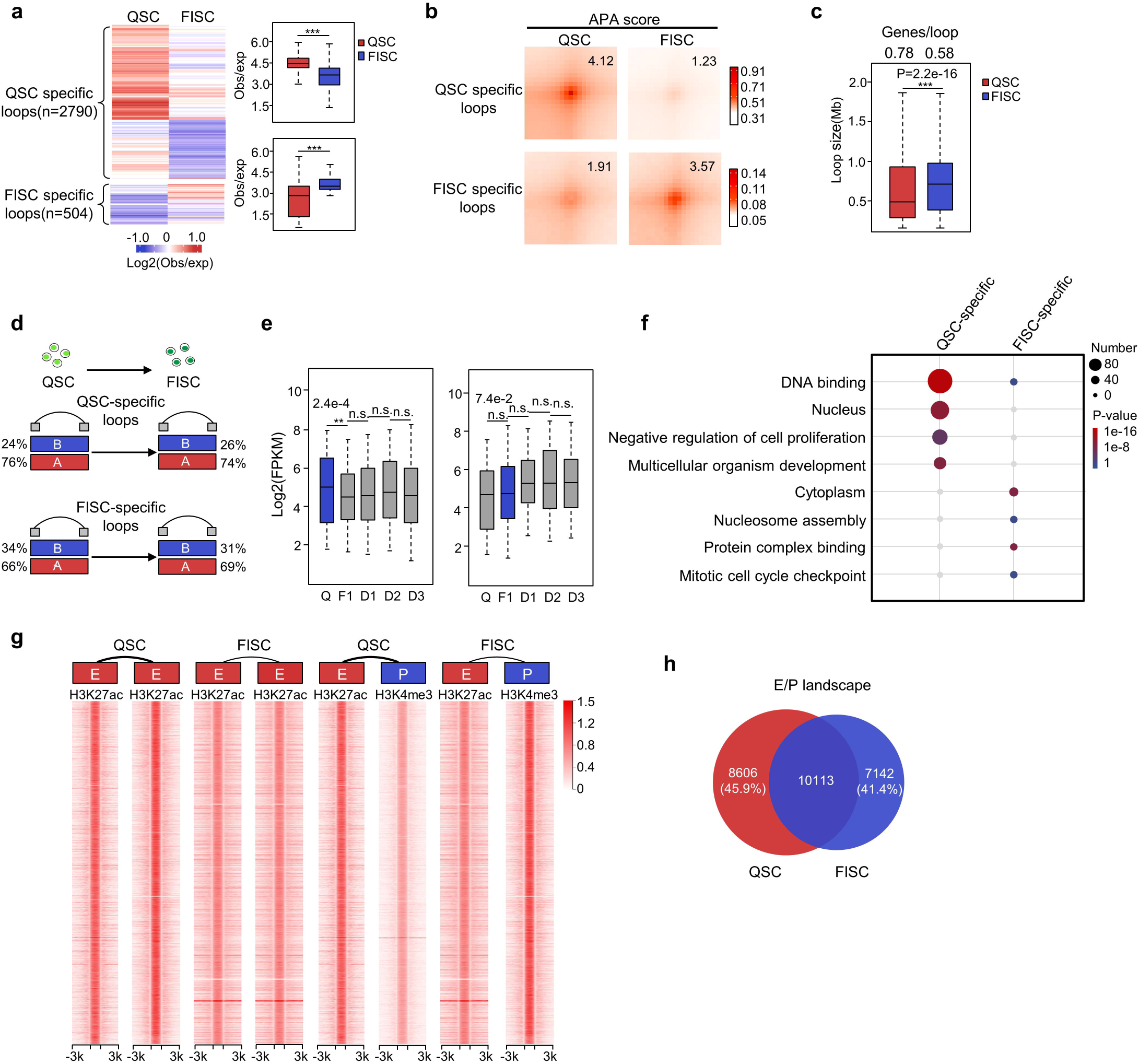

**Suppl. Fig. 6.**
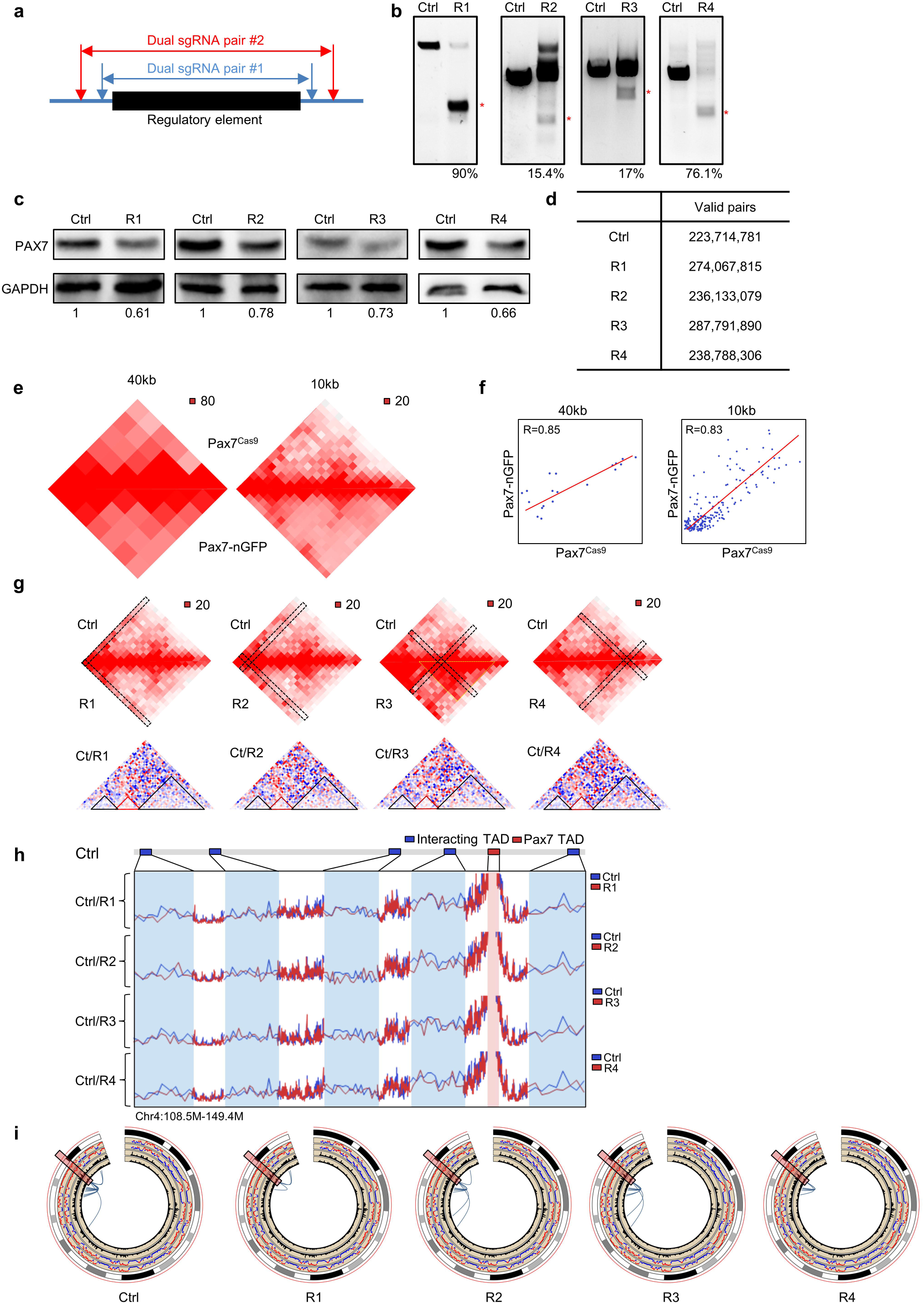

**Suppl. Fig. 7.**
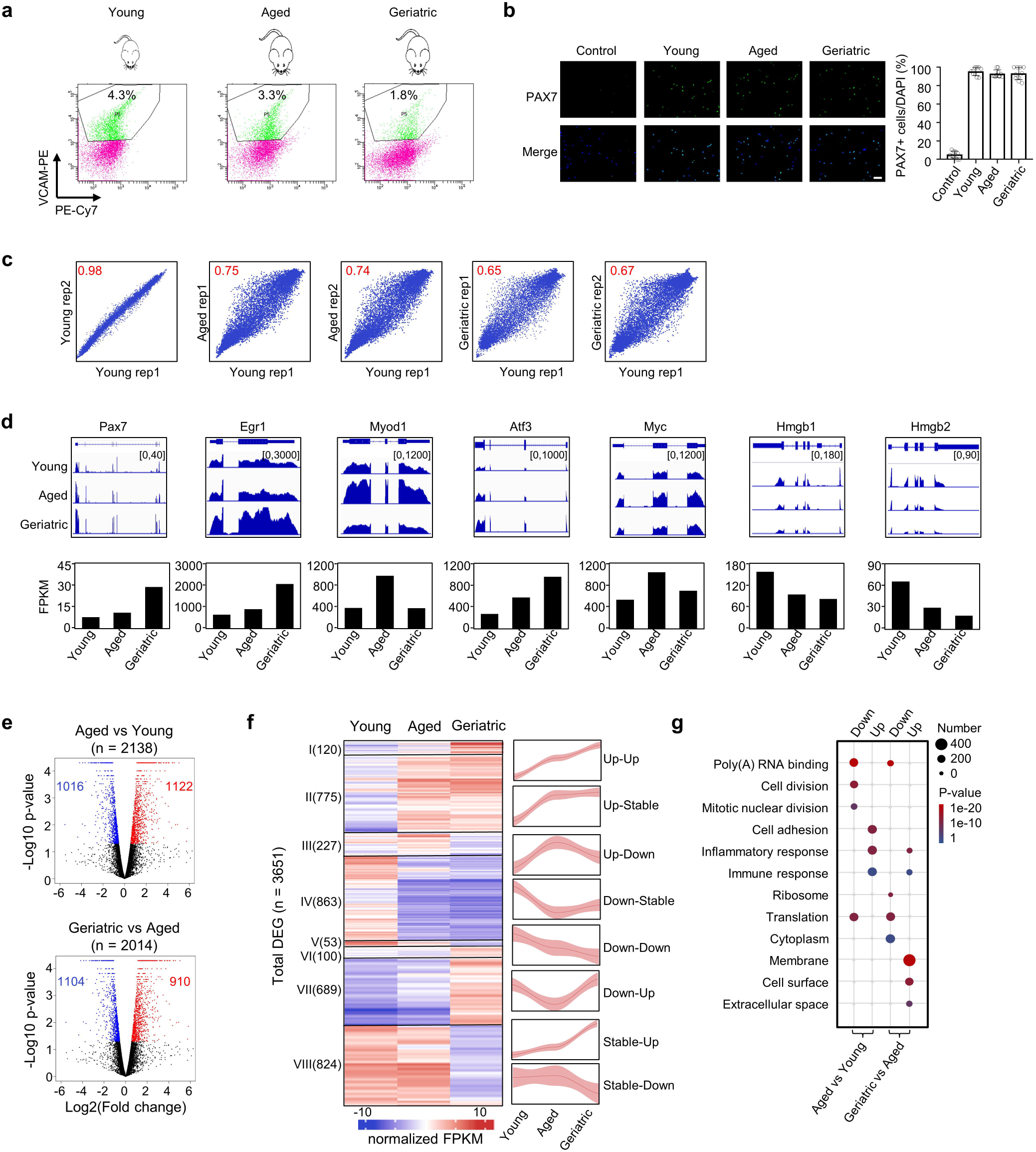

**Suppl. Fig. 8.**
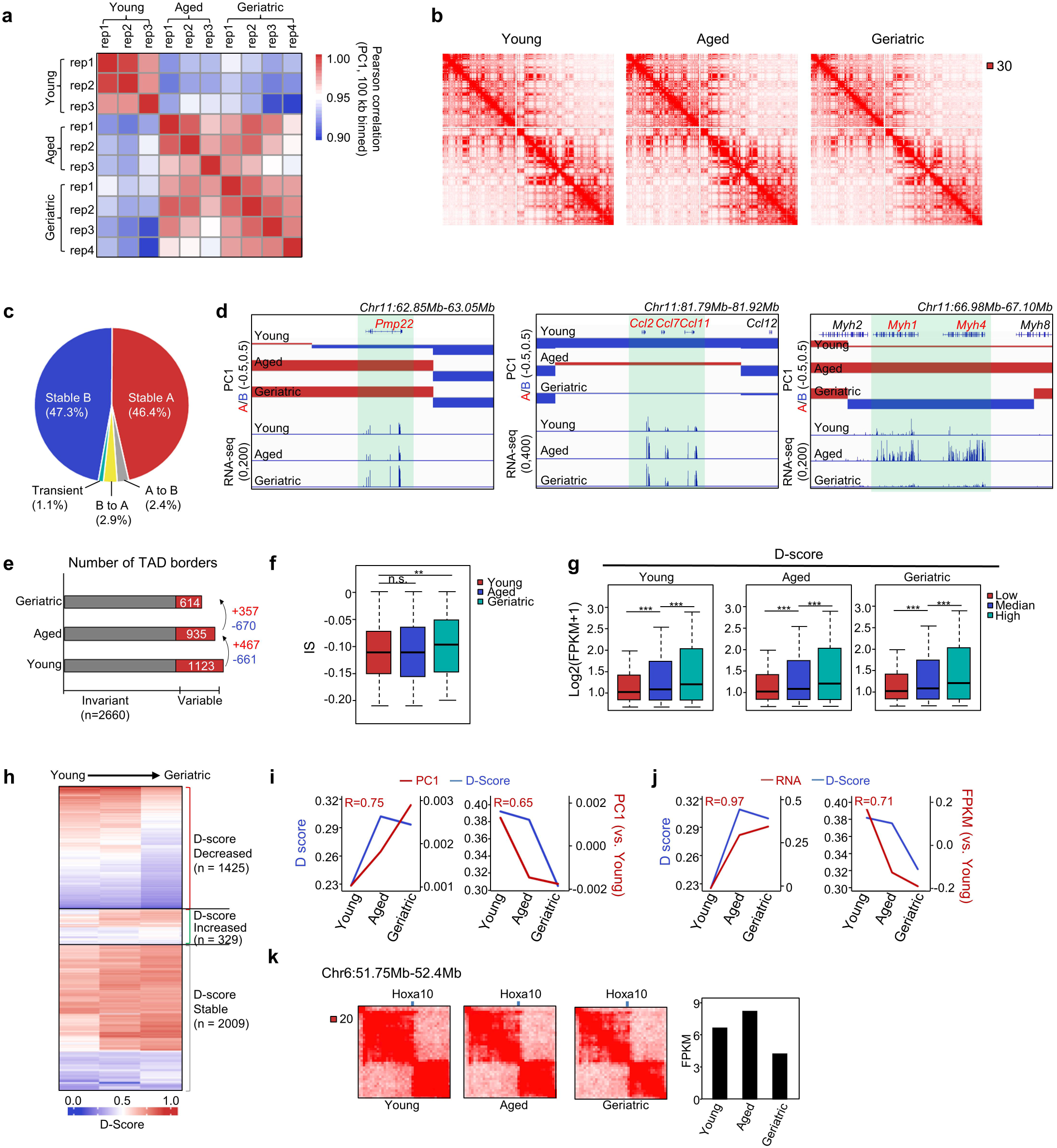

## Reference

1. Almada, A.E. & Wagers, A.J. Molecular circuitry of stem cell fate in skeletal muscle regeneration, ageing and disease. Nat Rev Mol Cell Biol 17, 267–279 (2016).

2. Yin, H., Price, F. & Rudnicki, M.A. Satellite cells and the muscle stem cell niche. Physiol Rev 93, 23–67 (2013).

3. Feige, P., Brun, C.E., Ritso, M. & Rudnicki, M.A. Orienting Muscle Stem Cells for Regeneration in Homeostasis, Aging, and Disease. Cell Stem Cell 23, 653–664 (2018).

4. Hwang, A.B. & Brack, A.S. Muscle Stem Cells and Aging. Curr Top Dev Biol 126, 299–322 (2018).

5. Bengal, E., Perdiguero, E., Serrano, A.L. & Munoz-Canoves, P. Rejuvenating stem cells to restore muscle regeneration in aging. F1000Res 6, 76 (2017).

6. Seale, P. et al. Pax7 is required for the specification of myogenic satellite cells. Cell 102, 777–786 (2000).

7. Bonev, B. et al. Multiscale 3D Genome Rewiring during Mouse Neural Development. Cell 171, 557–572 e524 (2017).

8. Rao, S.S. et al. A 3D map of the human genome at kilobase resolution reveals principles of chromatin looping. Cell 159, 1665–1680 (2014).

9. Stadhouders, R. et al. Transcription factors orchestrate dynamic interplay between genome topology and gene regulation during cell reprogramming. Nat Genet 50, 238–249 (2018).

10. Dixon, J.R. et al. Topological domains in mammalian genomes identified by analysis of chromatin interactions. Nature 485, 376–380 (2012).

11. Naumova, N. et al. Organization of the mitotic chromosome. Science 342, 948–953 (2013).

12. Guo, Y. et al. CRISPR Inversion of CTCF Sites Alters Genome Topology and Enhancer/Promoter Function. Cell 162, 900–910 (2015).

13. Kragesteen, B.K. et al. Dynamic 3D chromatin architecture contributes to enhancer specificity and limb morphogenesis. Nat Genet 50, 1463–1473 (2018).

14. Miguel-Escalada, I. et al. Human pancreatic islet three-dimensional chromatin architecture provides insights into the genetics of type 2 diabetes. Nat Genet 51, 1137–1148 (2019).

15. Paulsen, J. et al. Long-range interactions between topologically associating domains shape the four-dimensional genome during differentiation. Nat Genet 51, 835–843 (2019).

16. Beagrie, R.A. et al. Complex multi-enhancer contacts captured by genome architecture mapping. Nature 543, 519–524 (2017).

17. Quinodoz, S.A. et al. Higher-Order Inter-chromosomal Hubs Shape 3D Genome Organization in the Nucleus. Cell 174, 744–757 e724 (2018).

18. Du, Z. et al. Polycomb Group Proteins Regulate Chromatin Architecture in Mouse Oocytes and Early Embryos. Mol Cell 77, 825–839 e827 (2020).

19. Hug, C.B., Grimaldi, A.G., Kruse, K. & Vaquerizas, J.M. Chromatin Architecture Emerges during Zygotic Genome Activation Independent of Transcription. Cell 169, 216–228 e219 (2017).

20. Machado, L. et al. In Situ Fixation Redefines Quiescence and Early Activation of Skeletal Muscle Stem Cells. Cell Rep 21, 1982–1993 (2017).

21. van Velthoven, C.T.J., de Morree, A., Egner, I.M., Brett, J.O. & Rando, T.A. Transcriptional Profiling of Quiescent Muscle Stem Cells In Vivo. Cell Rep 21, 1994–2004 (2017).

22. Yue, L., Wan, R., Luan, S., Zeng, W. & Cheung, T.H. Dek Modulates Global Intron Retention during Muscle Stem Cells Quiescence Exit. Dev Cell 53, 661–676 e666 (2020).

23. Chandra, T. et al. Global reorganization of the nuclear landscape in senescent cells. Cell Rep 10, 471–483 (2015).

24. Zirkel, A. et al. HMGB2 Loss upon Senescence Entry Disrupts Genomic Organization and Induces CTCF Clustering across Cell Types. Mol Cell 70, 730–744 e736 (2018).

25. Sati, S. et al. 4D Genome Rewiring during Oncogene-Induced and Replicative Senescence. Mol Cell 78, 522–538 e529 (2020).

26. Sofiadis, K. et al. HMGB1 coordinates SASP-related chromatin folding and RNA homeostasis on the path to senescence. Mol Syst Biol 17, e9760 (2021).

27. Sousa-Victor, P. et al. Geriatric muscle stem cells switch reversible quiescence into senescence. Nature 506, 316–321 (2014).

28. Sousa-Victor, P., Garcia-Prat, L., Serrano, A.L., Perdiguero, E. & Munoz-Canoves, P. Muscle stem cell aging: regulation and rejuvenation. Trends Endocrinol Metab 26, 287–296 (2015).

29. Liu, L. et al. Chromatin modifications as determinants of muscle stem cell quiescence and chronological aging. Cell Rep 4, 189–204 (2013).

30. Schworer, S. et al. Epigenetic stress responses induce muscle stem-cell ageing by Hoxa9 developmental signals. Nature 540, 428–432 (2016).

31. Hernando-Herraez, I. et al. Ageing affects DNA methylation drift and transcriptional cell-to-cell variability in mouse muscle stem cells. Nat Commun 10, 4361 (2019).

32. Rocheteau, P., Gayraud-Morel, B., Siegl-Cachedenier, I., Blasco, M.A. & Tajbakhsh, S. A subpopulation of adult skeletal muscle stem cells retains all template DNA strands after cell division. Cell 148, 112–125 (2012).

33. Servant, N. et al. HiC-Pro: an optimized and flexible pipeline for Hi-C data processing. Genome Biol 16, 259 (2015).

34. Nagano, T. et al. Comparison of Hi-C results using in-solution versus in-nucleus ligation. Genome Biol 16, 175 (2015).

35. Rivas-Pardo, J.A. et al. Work Done by Titin Protein Folding Assists Muscle Contraction. Cell Rep 14, 1339–1347 (2016).

36. Linke, W.A. Titin Gene and Protein Functions in Passive and Active Muscle. Annu Rev Physiol 80, 389–411 (2018).

37. Shin, H. et al. TopDom: an efficient and deterministic method for identifying topological domains in genomes. Nucleic Acids Res 44, e70 (2016).

38. Wang, Q., Sun, Q., Czajkowsky, D.M. & Shao, Z. Sub-kb Hi-C in D. melanogaster reveals conserved characteristics of TADs between insect and mammalian cells. Nat Commun 9, 188 (2018).

39. Javierre, B.M. et al. Lineage-Specific Genome Architecture Links Enhancers and Non-coding Disease Variants to Target Gene Promoters. Cell 167, 1369–1384 e1319 (2016).

40. Li, L. et al. Widespread rearrangement of 3D chromatin organization underlies polycomb-mediated stress-induced silencing. Mol Cell 58, 216–231 (2015).

41. Novo, C.L. et al. Long-Range Enhancer Interactions Are Prevalent in Mouse Embryonic Stem Cells and Are Reorganized upon Pluripotent State Transition. Cell Rep 22, 2615–2627 (2018).

42. Kaul, A., Bhattacharyya, S. & Ay, F. Identifying statistically significant chromatin contacts from Hi-C data with FitHiC2. Nat Protoc 15, 991–1012 (2020).

43. Ancel, S., Stuelsatz, P. & Feige, J.N. Muscle Stem Cell Quiescence: Controlling Stemness by Staying Asleep. Trends Cell Biol 31, 556–568 (2021).

44. He, L., et al. CRISPR/Cas9/AAV9-mediated in vivo editing identifies MYC regulation of 3D genome in skeletal muscle stem cell. Stem Cell Reports (2021).

45. van Overbeek, M. et al. DNA Repair Profiling Reveals Nonrandom Outcomes at Cas9-Mediated Breaks. Mol Cell 63, 633–646 (2016).

46. Garcia-Prat, L. et al. FoxO maintains a genuine muscle stem-cell quiescent state until geriatric age. Nat Cell Biol 22, 1307–1318 (2020).

47. Evano, B. & Tajbakhsh, S. Skeletal muscle stem cells in comfort and stress. NPJ Regen Med 3, 24 (2018).

48. Zheng, H. et al. Resetting Epigenetic Memory by Reprogramming of Histone Modifications in Mammals. Mol Cell 63, 1066–1079 (2016).

49. Bian, Q., Anderson, E.C., Yang, Q. & Meyer, B.J. Histone H3K9 methylation promotes formation of genome compartments in Caenorhabditis elegans via chromosome compaction and perinuclear anchoring. Proc Natl Acad Sci U S A 117, 11459–11470 (2020).

50. Nora, E.P. et al. Targeted Degradation of CTCF Decouples Local Insulation of Chromosome Domains from Genomic Compartmentalization. Cell 169, 930–944 e922 (2017).

51. Fudenberg, G. et al. Formation of Chromosomal Domains by Loop Extrusion. Cell Rep 15, 2038–2049 (2016).

52. Heinz, S. et al. Transcription Elongation Can Affect Genome 3D Structure. Cell 174, 1522–1536 e1522 (2018).

53. Kim, Y.H. et al. Rev-erbalpha dynamically modulates chromatin looping to control circadian gene transcription. Science 359, 1274–1277 (2018).

54. Munoz-Canoves, P., Neves, J. & Sousa-Victor, P. Understanding muscle regenerative decline with aging: new approaches to bring back youthfulness to aged stem cells. FEBS J 287, 406–416 (2020).

55. Chen, W., Datzkiw, D. & Rudnicki, M.A. Satellite cells in ageing: use it or lose it. Open Biol 10, 200048 (2020).

56. Garcia-Prat, L. et al. Autophagy maintains stemness by preventing senescence. Nature 529, 37–42 (2016).

57. Sambasivan, R. et al. Pax7-expressing satellite cells are indispensable for adult skeletal muscle regeneration. Development 138, 3647–3656 (2011).

58. von Maltzahn, J., Jones, A.E., Parks, R.J. & Rudnicki, M.A. Pax7 is critical for the normal function of satellite cells in adult skeletal muscle. Proc Natl Acad Sci U S A 110, 16474–16479 (2013).

59. Liu, L., Cheung, T.H., Charville, G.W. & Rando, T.A. Isolation of skeletal muscle stem cells by fluorescence-activated cell sorting. Nat Protoc 10, 1612–1624 (2015).

60. Grieger, J.C., Choi, V.W. & Samulski, R.J. Production and characterization of adeno-associated viral vectors. Nat Protoc 1, 1412–1428 (2006).

61. Haeussler, M. et al. Evaluation of off-target and on-target scoring algorithms and integration into the guide RNA selection tool CRISPOR. Genome Biol 17, 148 (2016).

62. Chen, F. et al. YY1 regulates skeletal muscle regeneration through controlling metabolic reprogramming of satellite cells. EMBO J 38 (2019).

63. Zhao, Y. et al. MyoD induced enhancer RNA interacts with hnRNPL to activate target gene transcription during myogenic differentiation. Nat Commun 10, 5787 (2019).

64. Li, Y. et al. Long noncoding RNA SAM promotes myoblast proliferation through stabilizing Sugt1 and facilitating kinetochore assembly. Nat Commun 11, 2725 (2020).

65. Chen, X. et al. Translational control by DHX36 binding to 5’UTR G-quadruplex is essential for muscle stem-cell regenerative functions. Nat Commun 12, 5043 (2021).

66. Peng, X.L. et al. MyoD- and FoxO3-mediated hotspot interaction orchestrates super-enhancer activity during myogenic differentiation. Nucleic Acids Res 45, 8785–8805 (2017).

67. Huang, Y. et al. Large scale RNA-binding proteins/LncRNAs interaction analysis to uncover lncRNA nuclear localization mechanisms. Brief Bioinform 22 (2021).

68. Hnisz, D. et al. Super-enhancers in the control of cell identity and disease. Cell 155, 934–947 (2013).

69. Crane, E. et al. Condensin-driven remodelling of X chromosome topology during dosage compensation. Nature 523, 240–244 (2015).

70. Paulsen, J. et al. Chrom3D: three-dimensional genome modeling from Hi-C and nuclear lamin-genome contacts. Genome Biol 18, 21 (2017).

71. Stik, G. et al. CTCF is dispensable for immune cell transdifferentiation but facilitates an acute inflammatory response. Nat Genet 52, 655–661 (2020).

72. Di Stefano, M. et al. Transcriptional activation during cell reprogramming correlates with the formation of 3D open chromatin hubs. Nat Commun 11, 2564 (2020).

73. Serra, F. et al. Automatic analysis and 3D-modelling of Hi-C data using TADbit reveals structural features of the fly chromatin colors. PLoS Comput Biol 13, e1005665 (2017).

74. Pettersen, E.F. et al. UCSF Chimera--a visualization system for exploratory research and analysis. J Comput Chem 25, 1605–1612 (2004).

75. Wingender, E., Dietze, P., Karas, H. & Knuppel, R. TRANSFAC: a database on transcription factors and their DNA binding sites. Nucleic Acids Res 24, 238–241 (1996).

